# *APOE-*ε4-induced Fibronectin at the blood-brain barrier is a conserved pathological mediator of disrupted astrocyte-endothelia interaction in Alzheimer’s disease

**DOI:** 10.1101/2025.01.24.634732

**Authors:** Prabesh Bhattarai, Elanur Yilmaz, Elif Öykü Cakir, Hande Yüceer Korkmaz, Annie J. Lee, Yiyi Ma, Hilal Celikkaya, Mehmet Ilyas Cosacak, Verena Haage, Xue Wang, Nastasia Nelson, Weilin Lin, Yixin Zhang, Tal Nuriel, Dörthe Jülich, Özkan Iş, Scott A. Holley, Philip de Jager, Elizabeth Fisher, Kate Tubbesing, Andrew F. Teich, Taylor Bertucci, Sally Temple, Nilüfer Ertekin-Taner, Badri N. Vardarajan, Richard Mayeux, Caghan Kizil

## Abstract

Blood-brain barrier (BBB) dysfunction is a key feature of Alzheimer’s disease (AD), particularly in individuals carrying the *APOE-ε4* allele. This dysfunction worsens neuroinflammation and hinders the removal of toxic proteins, such as amyloid-beta (Aβ42), from the brain. In post-mortem brain tissues and in animal models, we previously reported that fibronectin accumulates at the BBB predominantly in *APOE-ε4* carriers. Furthermore, we found a loss-of-function variant in the fibronectin 1 (*FN1*) gene significantly reduces aggregated fibronectin levels and decreases AD risk among *APOE-ε4* carriers. Yet, the molecular mechanisms downstream of fibronectin at the BBB remain unclear. The extracellular matrix (ECM) plays a crucial role in maintaining BBB homeostasis and orchestrating the interactions between BBB cell types, including endothelia and astrocytes. Understanding the mechanisms affecting the ECM and BBB cell types will be critical for developing effective therapies against AD, especially among *APOE-ε4* carriers. Here, we demonstrate that *APOE-ε4*, Aβ42, and inflammation drive the induction of *FN1* expression in several models including zebrafish, mice, iPSC-derived human 3D astrocyte and 3D cerebrovascular cell cultures, and in human brains. Fibronectin accumulation disrupts astroglial-endothelial interactions and the signalling cascade between vascular endothelial growth factor (VEGF), heparin-binding epidermal growth factor (HBEGF) and Insulin-like growth factor 1 (IGF1). This accumulation of fibronectin in *APOE-ε4-* associated AD potentiates BBB dysfunction, which strongly implicates reducing fibronectin deposition as a potential therapeutic target for AD.

**Graphical abstract:** 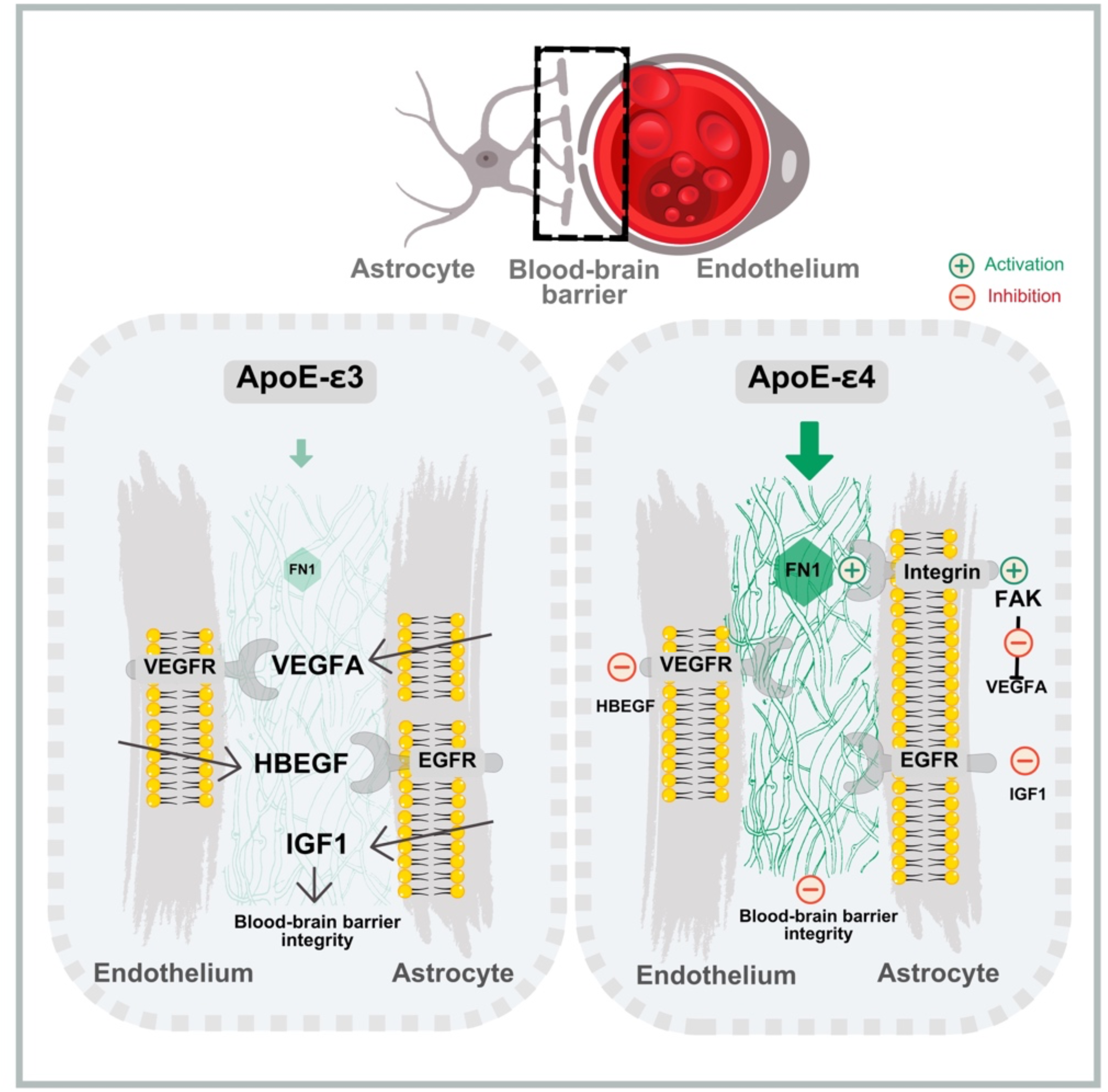

**Accessibility text:** This image illustrates the effects of different APOE isoforms (ApoE-ε3 and ApoE-ε4) on blood-brain barrier (BBB) integrity, focusing on the molecular interactions between astrocytes and endothelial cells. This figure emphasizes the detrimental effects of ApoE-ε4 on BBB integrity via fibronectin accumulation and altered signaling pathways.

The **top section** provides a schematic overview of the blood-brain barrier, highlighting astrocytes, endothelial cells, and their interface.

The **left panel** represents the ApoE-ε3 condition: Normal fibronectin (FN1) levels support healthy interactions between astrocytes and endothelial cells. Growth factors, including VEGFA, HBEGF, and IGF1, maintain BBB integrity through their respective receptors (VEGFR and EGFR). Green arrows indicate activation of these signaling pathways.

The **right panel** depicts the ApoE-ε4 condition: Elevated fibronectin (FN1) disrupts astrocyte-endothelium interactions. FN1 binds integrins and activates focal adhesion kinase (FAK), inhibiting VEGFA, which is required for endothelial HBEGF that in turn activates IGF1 signaling. Red symbols indicate inhibition of HBEGF, VEGFA, and IGF1 pathways, leading to BBB dysfunction.

**Highlights:** *APOE-ε4* drives fibronectin deposition in Alzheimer’s, disrupting astrocyte-endothelia interactions.

*APOE-ε4* and fibronectin co-localize, forming aggregates at blood-brain barrier (BBB).

Fibronectin alters the signaling between VEGF, IGF1, and HBEGF impairing BBB function.

Reducing fibronectin restores BBB integrity and offsets *APOE-ε4* pathology.

## Introduction

Impairments in the blood-brain barrier (BBB), a highly selectively coordinated network of cells that regulates the passage of substances between the bloodstream and the brain, has been well documented in Alzheimer’s disease (AD) (*1–7*). These alterations are exacerbated by the presence of the *APOE-ε4* allele, the strongest genetic risk factor for late-onset AD (*8–11*). A critical component of BBB dysfunction lies within the extracellular matrix (ECM). Aberrant remodeling of this dynamic three-dimensional network alters the vascular basement membrane architecture and contributes to AD pathogenesis (*2, 4, 12–15*). Specialized ECM components, such as collagen, laminin, and fibronectin, are instrumental in maintaining the tight balance between homeostatic neuronal functions, maintenance of the BBB, and anti-inflammatory activity of immune cells (*16, 17*). Thus, in AD, the interplay between ECM dysregulation and *APOE-ε4* is a critical mechanism driving BBB dysfunction.

We previously found that individuals with AD who carry one or more *APOE-ε4* alleles deposit significantly higher levels of fibronectin protein at the BBB (*18*). Intriguingly, we found that a loss-of-function variant in the *FN1* gene decreases fibronectin accumulation, reduces AD risk and delays the age at onset of disease in *APOE-ε4* homozygous carriers (*18*). These findings suggest fibronectin accumulation may be a downstream mediator of *APOE-ε4* in AD and implicates fibronectin-related pathways as potential therapeutic targets to mitigate BBB dysfunction and AD. However, the mechanisms by which fibronectin contributes to BBB dysfunction in AD remains largely unknown.

Here, we show that *APOE-ε4* and amyloid-β (Aβ42) induce the pathological accumulation of fibronectin at the BBB disrupting normal homeostatic astroglial-endothelial interactions by attenuating specific growth factor crosstalk. Using zebrafish and mouse models, 3D cultures of human iPSC-derived cerebrovascular cells, and human brain samples, we demonstrate that fibronectin deposition driven by *APOE-ε4* plays a key role in BBB dysfunction in AD pathogenesis.

## Results

### Fibronectin is induced in human brains with AD and vascular pathology

There is a strong association between AD and cerebrovascular pathology (CVP) (*5, 19–22*), but they can also exist independently (*23*). We hypothesized that AD and CVP may independently induce fibronectin deposition and contribute to BBB dysfunction. To test this hypothesis, we examined fibronectin upregulation as a shared pathological feature in AD and in CVP. We performed immunolabeling for fibronectin and CD31 (vascular endothelial marker) in dorsolateral prefrontal cortex (Brodmann area 9) of 18 postmortem human brains in four groups based on postmortem neuropathological assessments (**Fig. 1A-E**, **Table 1**): a) no CVP or AD (control brains; Braak stage 0-I, arteriolosclerosis score (ATS, indicating microvascular pathology) 0-1, n=2), b) CVP without AD (ATS 3-4, Braak stage 0-II, n=3), c) AD without CVP (Braak stage IV-VI, ATS 0-1, n=6), and d) coincident CVP and AD (Braak stage IV-VI, ATS 3-4, n=7) (**Table 1**). For these sets, we relied on the neuropathological observations because in *APOE-ε4* non-carriers, fibronectin is upregulated at the BBB with AD or CVP but exacerbated in *APOE-ε4* carriers (*18*). Fibronectin accumulation increased independently in group b (CVP: 62.9%, p = 1.9e-11), in group c (AD: 36.1%, p = 1.8e-10) and in group d (CVP+AD: 43.1%, p = 2.0e-11), when compared to control brains in group a (**Fig. 1G, H; Supplementary Table 1**), supporting the hypothesis that fibronectin deposition at the BBB is independently induced by AD and CVP.

**Fig. 1:**
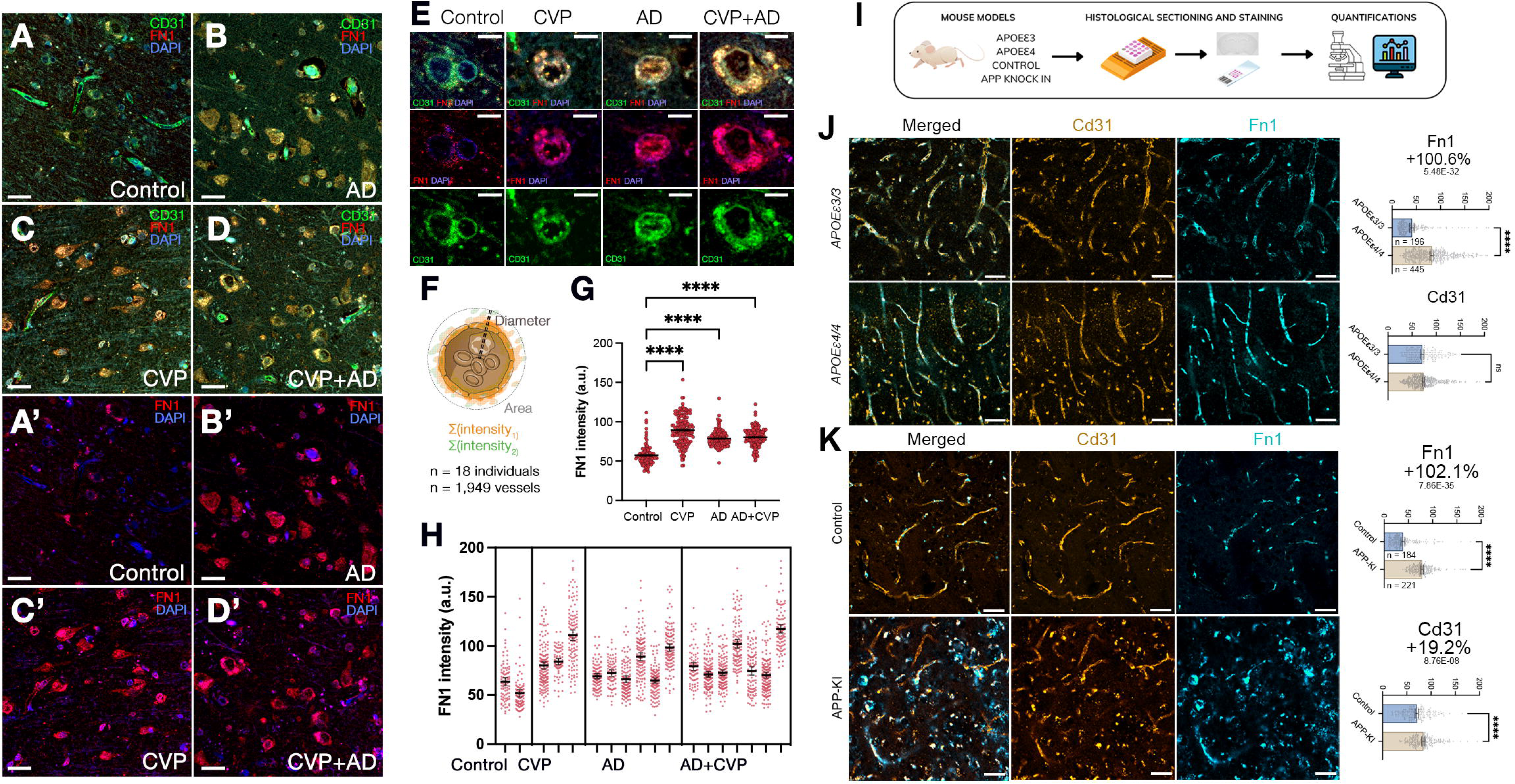
*APOE-ε4* induces Fibronectin in cerebrovascular pathology and Alzheimer’s disease. **A-E**, Double immunofluorescent staining (IFS) for CD31 (green) and Fibronectin 1 (FN1, red) with DAPI nuclear counterstain (blue) in control (**A, A’**), AD (**B, B’**), CVP (**C, C’**) and AD+CVP (**D, D’**) conditions. **E**, Higher magnification images from individual vessels are shown. **F**, Schematic depiction of the criteria for intensity and area measurements for 1,949 blood vessels. **G**, Mean intensity values for CD31 and FN1 in control, AD, CVP and AD+CVP conditions. **H,** Linear regression model showing intensity distribution for CD31, FN1 and DAPI in respect to vessel diameter. **I,** Schematic overview of the experimental approach illustrating the use of mouse models carrying *APOEε3/3*, *APOEε4/4* alleles, and an APP knock-in for AD pathology, the process of histological sectioning and staining, and subsequent quantification of results. **J,** Comparative analysis of FN1 expression in *APOEε3/3* and *APOEε4/4* mouse models. The left panel displays merged double IFS for CD31 (yellow) and Fibronectin 1 (FN1, cyan). Middle and right panels exhibit individual IFS for CD31 and FN1, respectively. Rightmost plots present quantifications of fluorescence intensity for FN1 and CD31 across a number of observations indicated as n, with corresponding percent changes and p-values. **K,** Assessment of FN1 expression in control and APP knock-in mouse model. Merged double IFS (left) and individual staining for Cd31 (middle) and Fn1 (right) are shown. The graphs quantify the fluorescence intensity for Fn1 and Cd31, providing statistical analysis with n representing the number of observations, as well as percent changes and p-values. Scale bars equal 50 µm (A-D’, J-K), 15 µm (E). Data is represented as dot plots or multivariable graphs, and statistical significance is noted as non-significant (ns) or with the precise p, t, F and W values where applicable with degrees of freedom indicated in parentheses. * (p < 0.0332), ** (p < 0.0021), *** (p < 0.002), and **** (p < 0.0001).

**Table 1:** Demographics of human brains used.

In order to assess the mechanism of fibronectin deposition more directly, we used knock-in mice that exhibit CVP (humanized targeted-replacement *APOE-ε4* and *APOE-ε3* mice (*24, 25*)) and amyloid proteinopathy (*APP* knock-in mice (*26*)). The use of the *APOE-ε4* and APP-knock-in mice allowed us to investigate the individual contributions of *APOE-ε4* and amyloid deposition to fibronectin induction. Humanized *APOE-ε4* mice show CVP (*27–29*), but not overt amyloid or tau pathology (*30–32*). APP-KI mice exhibit extensive amyloid pathology without overt vascular defects. We performed immunolabeling for fibronectin and Cd31 in both mouse strains followed by quantitative analyses from randomly selected regions of the cortex (**Fig. 1I-K**) to evaluate the contributions of *APOE-ε4* and amyloid deposition to fibronectin accumulation. In *APOE-ε4/4* mice, fibronectin accumulation was significantly higher compared to *APOE-ε3/3* mice (+100.6%, p = 5.46E-32), while Cd31 levels did not change (**Fig. 1J, Supplementary Table 1)**. Compared to non-transgenic age-matched control mice, APP-KI mice showed a significant increase in fibronectin (+102.1%, p=7.86E-35), as well as a slight increase in Cd31 (+19.2%, p=8.76E-08) (**Fig. 1K, Supplementary Table 1**). These results augment the observation in human brains that the role of fibronectin deposition is as a common pathological mediator of cerebrovascular dysfunction in AD (*2, 6, 7, 22, 33–36*).

### ApoE-ε4 induces FN1 in astrocytes and endothelia

Endothelia and astrocytes are prominent cell types of the BBB; thus, we hypothesized these cells contribute to fibronectin deposition in an ApoE-ε4-dependent manner. We used isogenic human iPSC-derived *APOE-ε3/3* and *APOE-ε4/4* astrocytes (*37*) to examine differences in fibronectin expression. *APOE-ε4/4* human astrocytes not only induced increased fibronectin expression but resulted in amorphic fibronectin deposition akin to pathological aggregates in humans in comparison to *APOE-ε3/3* astrocytes where fibronectin was deposited at lower levels and in fibrillar thread-like arrayed forms (**Fig. 2A, B**). Similarly, in iPSC-derived *APOE-ε3/3* and *APOE-ε4/4* endothelial cells, we also found elevated fibronectin expression in the *APOE-ε4/4* genotype (**Fig. 2C, D**). To determine whether fibronectin deposition around the blood vessels increased with *APOE-ε4/4* as we previously showed in human brain (*18*), we compared 3D vasculogenesis and angiogenesis model plexus (VAMP) cultures (*38*) containing isogenic iPSC-derived endothelial and mural cells with *APOE-ε3/3* and *APOE-ε4/4* genotypes. We found that compared to *APOE-ε3/3*, *APOE-ε4/4* leads to increased deposition proximal to the CD31-positive vascular structures (**Fig. 2E**). This suggests that *APOE-ε4*-dependent fibronectin production by both astrocytes and endothelia contributes to pathological accumulation at the BBB.

**Fig. 2:**
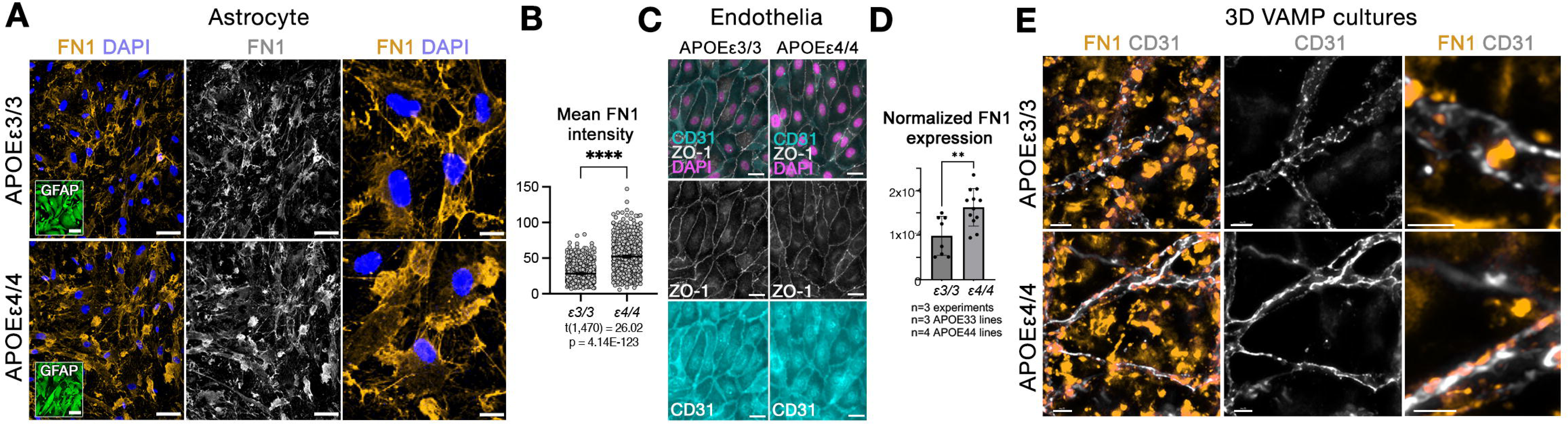
*APOE-ε4* induces Fibronectin in endothelia and astrocytes. **A**, Immunofluorescence staining (IFS) for FN1 (yellow) with DAPI counterstain (blue) in APOEε3/3 and APOEε4/4 isogenic iPSC-derived human astrocytes. Single channel in white shows FN1. Rightmost panel is high magnification on FN1 deposition. **B,** Quantification of FN1 intensity in single cells and their close periphery. **C,** Human iPSC-derived APOEε3/3 and APOEε4/4 endothelial cultures showing endothelial markers ZO-1 (white) and CD31 (cyan) with DAPI counterstain (violet). **D,** Quantitative real-time PCR results in APOEε3/3 and APOEε4/4 endothelia for FN1 gene expression. **E,** 3D VAMP cultures from APOEε3/3 and APOEε4/4 iPSCs stained for CD31 (white) and FN1 (yellow), showing FN1 deposition at the vascular interface. Scale bars equal 25 µm (A, high magnification image: 10 µm), 20 µm (C), 200 µm (E). Data is represented as dot plots or multivariable graphs, and statistical significance is noted as non-significant (ns) or with the precise p, t, F and W values where applicable with degrees of freedom indicated in parentheses. * (p < 0.0332), ** (p < 0.0021), *** (p < 0.002), and **** (p < 0.0001).

### ApoE-ε4 and fibronectin co-localize

ApoE is known to form extracellular aggregates (*39–41*), and we found fibronectin deposition also forms extracellular deposits (*18*). In *APOE-ε4/4* astrocytes, fibronectin deposits showed large aggregate structures that co-localize with ApoEε4, whereas in *APOE-ε3/3* astrocytes, no such aggregates were observed. There was also increased ApoE intensity in *APOE-ε4/4* astrocytes (**Fig. 3A,B**). In individual cells in culture, we observed that in *APOE-ε4/4* astrocytes, at any given ApoE protein intensity, there was a consistent increase in the amount of fibronectin in comparison to *APOE-ε3/3* astrocytes. In *APOE-ε4/4* astrocytes both ApoE and fibronectin levels were significantly higher than in *APOE-ε3/3* astrocytes (**Fig. 3C**, ε3: *R^2^*=0.049, ε4: *R^2^*=0.396). Co-localization in *APOE-ε4/4* astrocytes show a greater overlap of ApoE and fibronectin extracellular protein deposits (**Fig. 3D**). This implies that fibronectin might also co-aggregate with ApoEε4. Immunolabeling of ApoE, fibronectin and glial fibrillary acidic protein (GFAP) in post-mortem AD brains (n=2 *APOE-ε4/*4 and n=2 *APOE-ε3/3*) (**Fig. 3E,F**), confirmed our *in vitro* findings. We found that fibronectin and ApoE depositions significantly increase at the BBB encircled by astrocytic endfeet (**Fig. 3F,G)** in *APOE-ε4/4*-associated AD in comparison to *APOE-ε3/3* AD. The correlation of fibronectin and ApoE deposition was also significantly increased in *APOE-ε4* carriers compared to non-carriers (**Fig. 3H**, ε3: *R^2^*=0.108, ε4: *R^2^*=0.154).

**Fig. 3:**
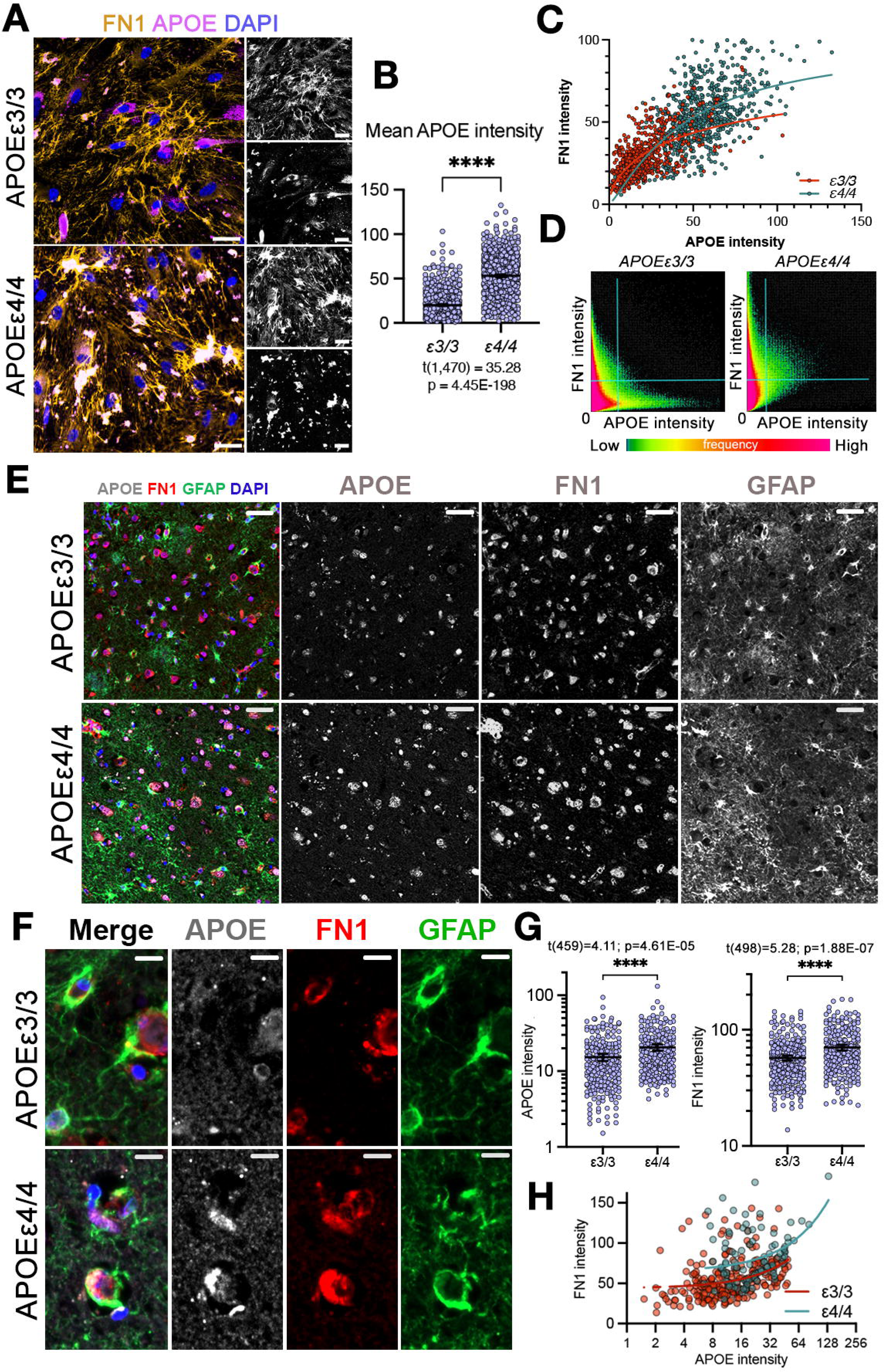
APOEε4 and FN1 co-localize with APOEε4. **A,** Immunofluorescence staining (IFS) for FN1 (yellow), APOE (purple) with DAPI counterstain (blue) in APOEε3/3 and APOEε4/4 isogenic iPSC-derived human astrocytes. Single channels in white shows FN1 and APOE. **B,** Quantification of APOE protein intensity in extracellular locations. **C,** Correlation graph for APOE and FN1 intensity in APOEε3/3 and APOEε4/4 expressing astrocytes. **D,** Co-localization histogram for APOE and FN1. Upper right quadrant shows co-localization and is enriched in APOEε4/4. **E**, IFS for FN1 (red), APOE (white), and GFAP (green) with DAPI counterstain (blue) in APOEε3/3 and APOEε4/4 post-mortem human brain samples of AD cases. Single channels in white shows FN1, APOE and GFAP channels alone. **F**, High magnification views from a smaller region of brains in E, showing co-localization of APOE and FN1 at GFAP-enwrapped BBB predominantly in APOEε4/4 case. **G,** Quantification of APOE and FN1 intensities in human brains. **H,** Correlation graph for APOE and FN1 intensity in APOEε3/3 and APOEε4/4 AD cases. **I,** Schematic model for mechanism of the pathogenicity of FN1 at the blood-brain barrier. 25 µm (A), 50 µm (E), 15 µm (F). Data is represented as dot plots or multivariable graphs, and statistical significance is noted as non-significant (ns) or with the precise p and t values where applicable with degrees of freedom indicated in parentheses. * (p < 0.0332), ** (p < 0.0021), *** (p < 0.002), and **** (p < 0.0001).

### Inflammation enhances FN1 deposition

ApoEε4 and brain amyloidosis amplify inflammation (*29, 42, 43*), and are linked to ECM remodeling during tissue repair (*12, 44, 45*). We used primary human astroglia (pHA) cultures (*46–51*) and starPEG-heparin-based hydrogel system to model inflammation and mimic brain tissue mechanics and biochemical interactions *in vitro* (*46, 49, 50*). Embedding pHA in 3D hydrogels recapitulated key brain microenvironment features, including fibronectin- and laminin-rich ECM structures and astroglia-neuron networks (*50*). Treatment with TNFα, a systemic inflammation mediator, induced reactive astrocytic states and NFκB signaling (**Fig. 4, Supplementary Figs. 1-3**), upregulating FN1 expression. The increased expression of FN1 was reversed after co-treatment with IL4, an anti-inflammatory cytokine that attenuates gliosis and modulates TNFα-driven changes (*50, 52, 53*) (**Fig. 4A-C**). Since pHAs express TNFα and IL4 receptors (*50*) (**Supplementary Fig. 3A**), we hypothesized direct cytokine regulation of astroglia. Comparing our dataset with prior analyses of 3D pHA cultures treated with Aβ42 (amyloid toxicity) and KYNA (excitotoxicity) (*46, 50*), we found TNFα significantly upregulated *FN1* gene expression, while IL4 reduced fibronectin levels alone or with TNFα or Aβ42 (**Fig. 4D).** Similarly, iPSC-derived endothelial cells also respond to inflammatory cytokines TNFα and IL1β by increased FN1 expression (**Supplementary Fig. 4B**), suggesting a mechanistic link between inflammation and fibronectin expression in astrocytes and endothelia. We validated these findings in AD brains using single-nucleus RNA sequencing of 1.64 million nuclei from 424 samples from the dorsolateral prefrontal cortex in the Religious Orders Study and the Memory and Aging Project (ROS/MAP) (*54*). Fibronectin was linked to pathological AD, particularly astrocyte cluster 5 (b: 0.0300, se: 0.0057, FDR p: 1.55E-06), characterized by inflammation, gliosis, and expression of *SERPINA3* (**Fig. 4E),** a reactive astrocyte marker (*54, 55*) that is also upregulated in TNFα-treated 3D astrocytes (**Fig. 4C**). Immunolabeling post-mortem brains confirmed fibronectin upregulation in reactive astrocytes (**Fig. 4F**, n=4). We validated this observation *in vivo* using zebrafish model where fibronectin induction by TNFα elevated NFκB signaling and fibronectin deposition around astrocytes, correlating with NFκB activity (**Fig. 4G-I, Supplementary Fig. 4**). Camptothecin and Torin2, which we previously found to promote anti-inflammatory states by modulating microglial polarization *in vitro* and *in vivo* (*56*), rescued amyloid-induced fibronectin deposition and astrocytic hypertrophy at the BBB in a zebrafish model of Aβ42-induced inflammation and fibronectin deposition (*18, 20, 47, 48, 52, 56–67*), marked by astrocytic Glutamine synthetase levels (**Fig. 4J-K, Supplementary Table 1**). These findings demonstrate that inflammation amplifies fibronectin deposition, consistent with the evidence that TNFα drives fibronectin-induced astrocyte reactivity (*14, 68*).

**Fig. 4:**
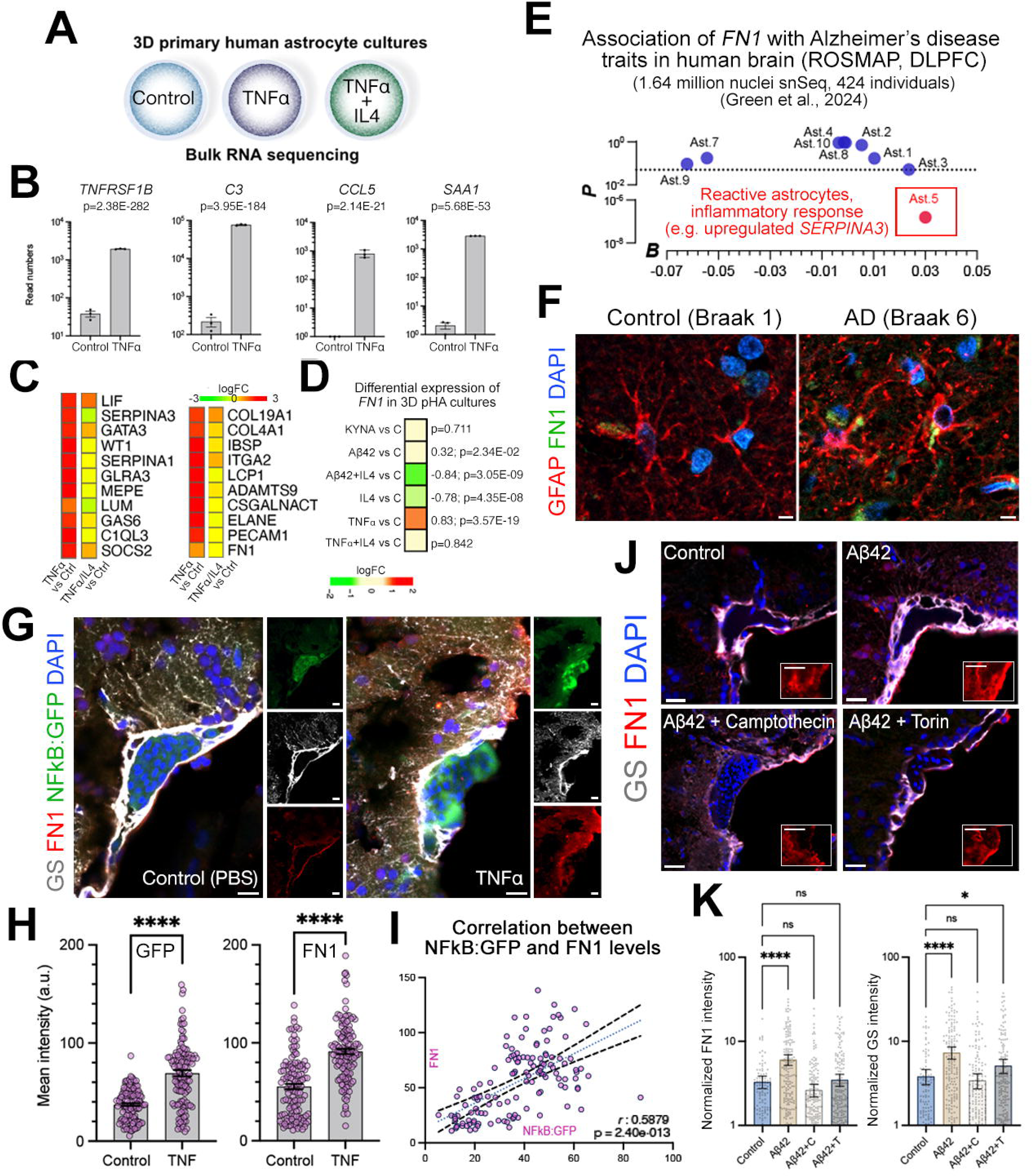
Inflammation induces Fibronectin in reactive astrocytes. **A,** Schematics for culture and treatment conditions for bulk RNA sequencing in 3D primary human astrocytes (pHA) cultures. **B**, RNA read numbers for selected TNFα downstream genes *TNFRSF1B, C3, CCL5* and *SAA1* in TNFα-treated pHA cultures. **C**, Heatmap for differential expression of selected TNFα-targets and reactive astrocyte markers in TNFα vs control and TNFα+IL4 vs control 3D pHA cultures. **D**, Heatmap of differential expression of *FN1* in several toxicity conditions in 3D pHA cultures (KYNA: Kynurenic acid, Aβ42: amyloid beta42). KYNA, Aβ42 and Aβ42+IL4 from ref.(*50*). **E**, Analyses of association of *FN1* with human AD traits in ROSMAP cohort astrocyte clusters positively associate *FN1* with reactive astrocyte cluster 5. **F,** Immunostaining for GFAP (red) and FN1 (green) with DAPI (blue) counterstain in control and AD postmortem human brain. FN1 is detected in hypertrophic reactive astrocytes. **G,** IFS for GS (white), FN1 (red) and NFkB:GFP (green) in control and TNFα injected zebrafish brains. Smaller panels show individual color-coded channels. **H**, Quantification of GFP and FN1 intensities. **I**, Interaction graph for NFkB:GFP and FN1 levels. **J**, Immunostaining for GS (white) and FN1 (red) with DAPI (blue) counterstain in control, Aβ42 injected, Aβ42+Camptothecin-injected, and Aβ42+Torin-injected zebrafish brains. Images highlight dorsoventral vascular hub. Insets: FN1 channel alone. **K**, Quantification graph for FN1 and GS intensities normalized to DAPI. Every data point is independent gliovascular contact point. Scale bars equal 10 µm (**F**) and 25 µm (**G, J**). Quantitative data in the graphs are represented as dot plots, and statistical significance is noted as non-significant (ns) or with the precise p-value where applicable. * (p < 0.0332), ** (p < 0.0021), *** (p < 0.002), and **** (p < 0.0001).

### Fibronectin disrupts BBB integrity via attenuated VEGF/HBEGF/IGF1 signaling cascade

To investigate the potential downstream effects of increased fibronectin, we performed single cell transcriptomics in *fn1b* knockout zebrafish (*fn1b*^-/-^) (*69*), which blocks the upregulation of fibronectin after Aβ42 exposure (*18*). We performed differential gene expression and KEGG pathways analyses across 32,294 cells in 38 clusters by comparing cells from wild-type (+Aβ42) and *fn1b*^-/-^ (+Aβ42) zebrafish (**Fig. 5A-C, Supplementary Fig. 5, Supplementary Table 5**). We identified significant enrichment of VEGF (p=3.8E-02, OR=2.87) and Insulin pathways (p=4.3E-03, OR=35.78), mediated by MAPK signaling (p=4.0E-05, OR=3.41), and a reduction in ECM-receptor interaction pathways (p=5.9E-04, OR=0.15) (**Fig. 5D,E**). These findings show that fibronectin modulates VEGF and Insulin signaling by altering ECM activity. Since VEGF signalling - essential for BBB integrity and vascular-astrocytic communication - is disrupted in AD, leading to BBB breakdown (*1, 57, 70–74*), we hypothesized that reduced VEGF signaling is a mediator of the pathological effects of fibronectin. Astrocytes in *fn1b* knockout zebrafish upregulated NFκB inhibitor *nfkbib* (*75*), VEGF ligand *vegfa*, insulin growth factor 1 *igf1*, and VEGF signaling inducer *irfbp2b* (*76*), while *fn1b* knockout endothelia upregulated NFκB-related genes that potentiate VEGF signaling (*77, 78*) (**Fig. 5F-G, Supplementary Table 5**). These findings align with human AD brains, where single-nucleus transcriptomics showed astrocytic VEGFA reductions across disease stages (**Supplementary Fig. 6**) (*2, 33, 79–83*), accompanied by decreased VEGFA protein in post-mortem AD samples and *APOE-ε4/4* iPSC-derived astrocytes (**Fig. 5H-J**). Since fibronectin binds integrins (*14, 84*) that are expressed on astrocytes (**Supplementary Fig. 7**), tested the hypothesis that inhibition of Focal Adhesion Kinase (FAK) – intracellular mediator of integrin signal cascade (*85*) - could restore VEGFA expression by treating zebrafish with the selective FAK inhibitor PF573228 (*85*). This restored VEGFA expression under Aβ42 conditions (**Fig. 5K**). To investigate the downstream mechanisms of the FN1/VEGF signaling axis, we used double-transgenic zebrafish (Tg(fli1a:EGFP)/Tg(her4.1:RFP) (*86, 87*) marking cerebrovascular endothelia (green) and astrocytes (red) with fluorescent reporters, and inhibited VEGF signaling with VEGFR inhibitors (Tivozanib, ZM306416, and Semaxanib). Single-cell transcriptomics of 22,712 sorted cells (17% endothelia, 39.2% astrocytes) identified 1,557 differentially expressed genes (DEGs) (**Fig. 6A-C, Supplementary Fig. 8, Supplementary Table 6**). To find the potential genes proximally regulated by VEGF signalling, we analyzed previously documented VEGF target genes (*88, 89*) among the DEGs, and found that VEGFR inhibition significantly downregulated endothelial *hbegf* isoforms, validated in kdrl:GFP zebrafish, a fluorescent reporter for endothelia (**Fig. 6E-F**). This is consistent with the upregulation of *hbegf* in *fn1b^-/-^*knockout validated by scSeq and tissue immunolabeling (**Fig. 6G**) and indicates that reduced endothelial HBEGF downstream of FN1/VEGF signaling also contributes to BBB pathology.

**Fig. 5:**
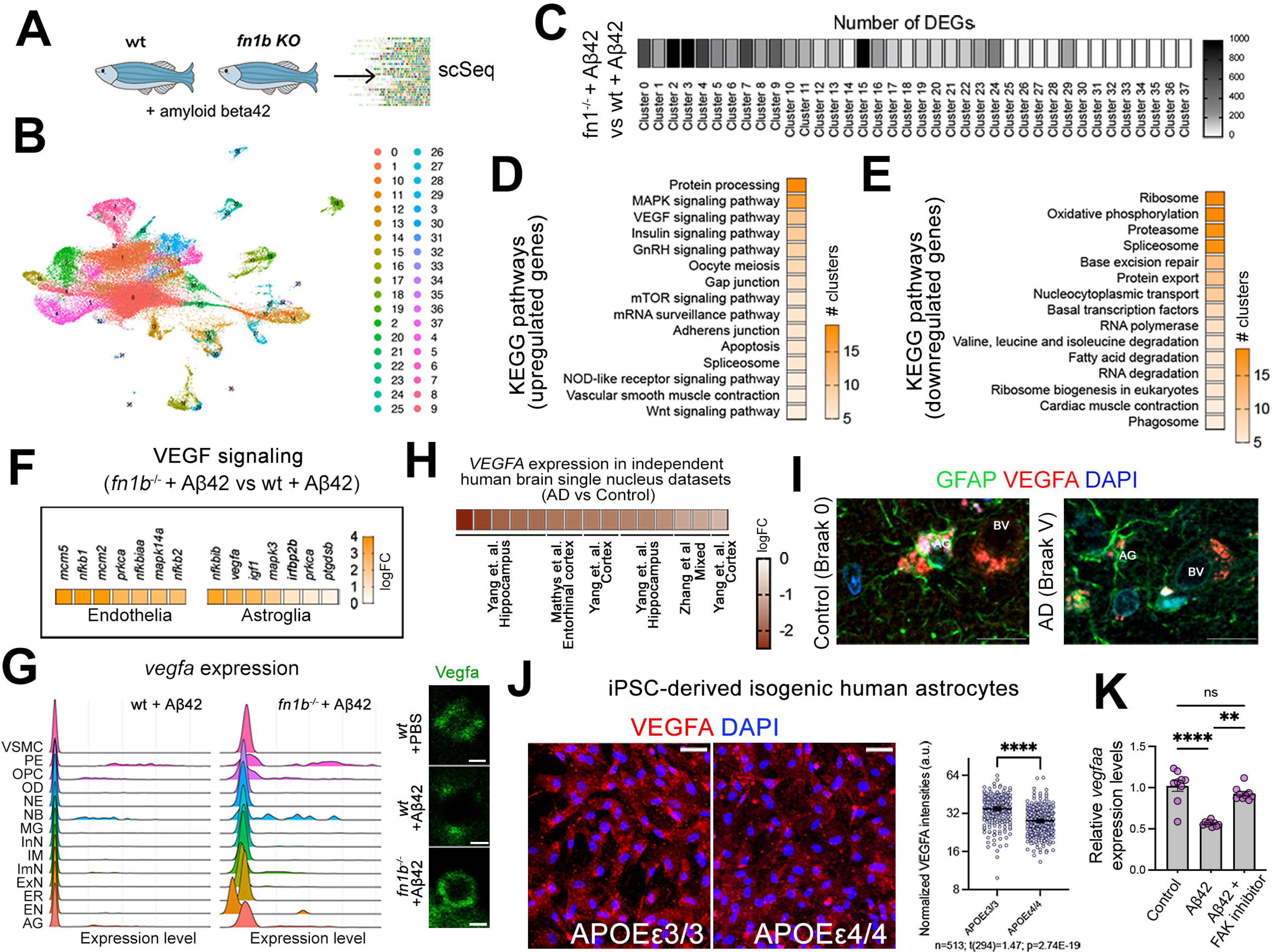
Fibronectin suppresses vascular endothelial growth factor signaling. **A,** Schematics for single cell sequencing comparison in wild-type control and *fn1b* homozygous knockout (*fn1b*^-/-^) both after Aβ42. **B**, UMAP plot for 38 cell clusters identified. **C**, Heat map showing the number of differentially expressed genes in every cluster. **D, E**, Heat map showing the number of times a certain KEGG pathway identifier is enriched in identified clusters. Upregulated genes (**D**) and downregulated genes (**E**) were analyzed separately depicted with separate heat maps. **F**, Selected top differentially expressed genes within the enriched VEGF signaling related KEGG pathway in endothelia and astrocytes. **G**, Ridge plots of *vegfa* expression and representative VEGFA IFS in wild type + Aβ42 and *fn1b*^-/-^ + Aβ42 zebrafish brains. Astrocytes increase *vegfa* expression upon *fn1b* loss-of-function. (AG: astrocytes, VSMC: vascular smooth muscle cell, OPC: oligodendrocyte precursor cell, PE: pericyte, OD: oligodendrocyte, NB: neuroblast, NE: neuron, MG: microglia, InN: inhibitory neurons, ExN: excitatory neuron, IM: immune cell, ImN: immature neuron, ER: erythroid cells, EN: endothelia). **H**, *VEGFA* expression changes in publicly available human single nucleus datasets from control and AD human brains (from refs.(*2, 33, 79*)). **I** IFS for VEGFA and GFAP with DAPI counterstain in control and AD post-mortem human brain show reduced VEGFA in astrocytes (AG) and around the blood vessels (BV). **J**, VEGFA protein in APOEε3/3 and APOEε4/4 expressing iPSC-derived human astrocytes, and quantification of VEGFA protein intensities. **K**, Quantitative real time PCR for *vegfaa* in control, Aβ42 -injected, Aβ42 + FAK inhibitor-injected zebrafish brains. Scale bars equal 5 µm (**G**) or 25 µm (**I, J**). Data is represented as dot plots or multivariable graphs, and statistical significance is noted as non-significant (ns) or with the precise p, t, F and W values where applicable with degrees of freedom indicated in parentheses. * (p < 0.0332), ** (p < 0.0021), *** (p < 0.002), and **** (p < 0.0001).

**Fig. 6:**
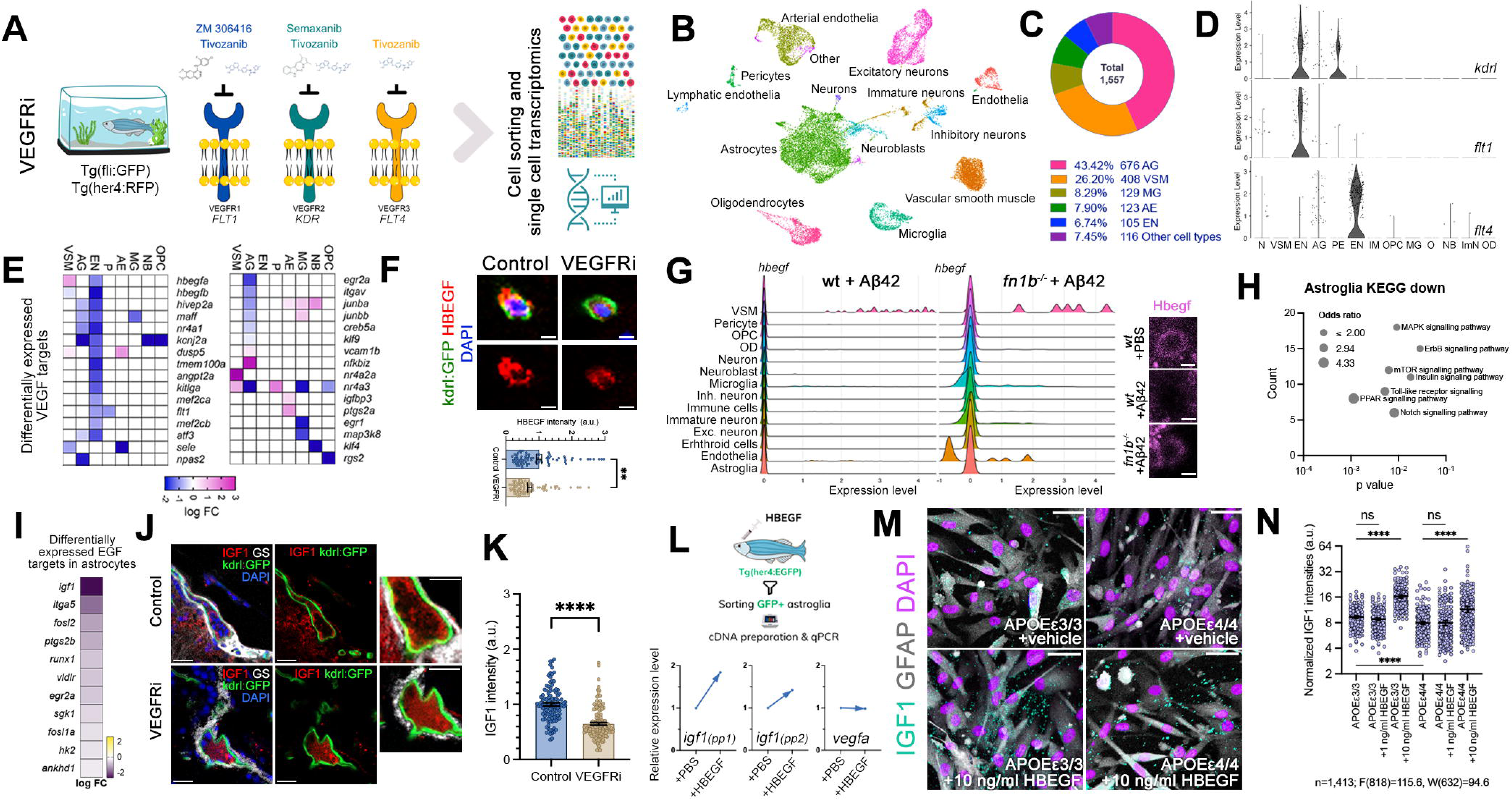
VEGF signaling is required for HBEGF that induces IGF1. **A**, Schematic view of VEGF receptor inhibition study (VEGFRi) and subsequent single cell transcriptomics in double reporter zebrafish line Tg(fli1a:EGFP) marking endothelia and Tg(her4:RFP) marking astrocytes. **B**, UMAP for identified cell types. **C**, Distribution of differentially expressed genes to cell types. **D**, Violin plots for expression of VEGF receptors *kdrl*, *flt1* and *flt4* in cell types (AG: astrocytes, VSM: vascular smooth muscle, OPC: oligodendrocyte precursor cell, PE: pericyte, OD: oligodendrocyte, NB: neuroblast, N: neuron, MG: microglia, IM: immune cell, ImN: immature neuron, EN: endothelia, O: Other). **E**, Heat map of differential expression of selected VEGF target genes in major cell types from (B) and (C). **F**, Immunostaining for HBEGF (red) and kdrl:GFP (green) with DAPI counterstain (blue) in control and VEGFRi zebrafish brains showing single vasculature, and quantification graph for HBEGF. VEGFRi reduces HBEGF levels. **G**, Ridge plot for *hbegfb* expression in control + Aβ42 and *fn1b*^-/-^ + Aβ42 zebrafish brains indicating upregulation in endothelia. **H**, Multiple variable graph for downregulated pathways in astrocytes with VEGFRi. MAPK, ErbB, mTOR, Insulin signaling pathways are enriched. **I**, Heat map for differential expression of selected EGF target genes in astrocytes after VEGFRi. **J**, IFS for IGF1 (red), GS (white), kdrl:GFP (green) with DAPI counterstain (blue) in control and VEGFRi zebrafish brains. **K**, Quantification of IGF1 levels. Every data point is a gliovascular contact point. **L**, Quantitative real time PCR setup where sorting her4:EGFP-positive astrocytes after HBEGF injection, which increases *igf1* expression determined by two primer pairs (pp) while *vegfa* expression is unchanged in adult zebrafish brain (reference gene: *b-actin*). **M,** Immunocytochemistry for IGF1 (cyan) and GFAP (white) with DAPI counterstain (violet) in APOEε3/3 and APOEε4/4 expressing isogenic astrocytes with and without 10 ng/ml HBEGF treatment. **N,** Quantification of IGF1 intensities normalized to cell numbers. Scale bars equal 5 µm (**F, G**), 25 µm (**J, M**). Data is represented as dot plots or multivariable graphs, and statistical significance is noted as non-significant (ns) or with the precise p, F and W values where applicable with degrees of freedom indicated in parentheses. * (p < 0.0332), ** (p < 0.0021), *** (p < 0.002), and **** (p < 0.0001).

HBEGF, binding to the EGF receptor (ERBB1), regulates inflammation, neurogenesis, and astrocyte maturation via MAPK and ERBB1 signaling (*90–94*). Thus, fibronectin-dependent reduction of HBEGF could alter astrocytes in the gliovascular niche. VEGFR inhibition (VEGFRi) in zebrafish revealed downregulated MAPK, ERBB, and Insulin pathways, reflecting changes observed in *fn1b*^-/-^ knockout (**Fig. 5D**, **Fig. 5H**). Notably, known EGF targets (*95–98*), including *igf1*, were significantly reduced (**Fig. 6I, Supplementary Table 6**), a finding further confirmed by immunolabeling (**Fig. 6J-K**). HBEGF induced *igf1* expression without affecting *vegfa* levels (**Fig. 6L**), and in human iPSC-derived astrocytes, *APOE-ε4* reduced IGF1 levels, which were restored by HBEGF in a dose-dependent manner (**Fig. 6M-N**), indicating that HBEGF is upstream of IGF1.

Since IGF1 is crucial for BBB function (*99–104*), these findings indicate that FN1/VEGF/HBEGF/IGF1 signaling axis regulates gliovascular homeostasis, which is disrupted in AD via *APOE-ε4*-dependent elevated fibronectin deposition. To assess if HBEGF and IGF1 could reverse fibronectin-mediated VEGF reduction and BBB damage, we analyzed gliovascular structure and tight junctions in VEGFRi-treated zebrafish with or without ectopic HBEGF/IGF1. VEGFRi compromised endothelial integrity as ZO-1/GFP colocalization was significantly reduced (**Fig. 7A-B**), consistent with previous findings (*57, 71*), while HBEGF/IGF1 treatment restored tight junction levels (**Fig. 7B**).

**Fig. 7:**
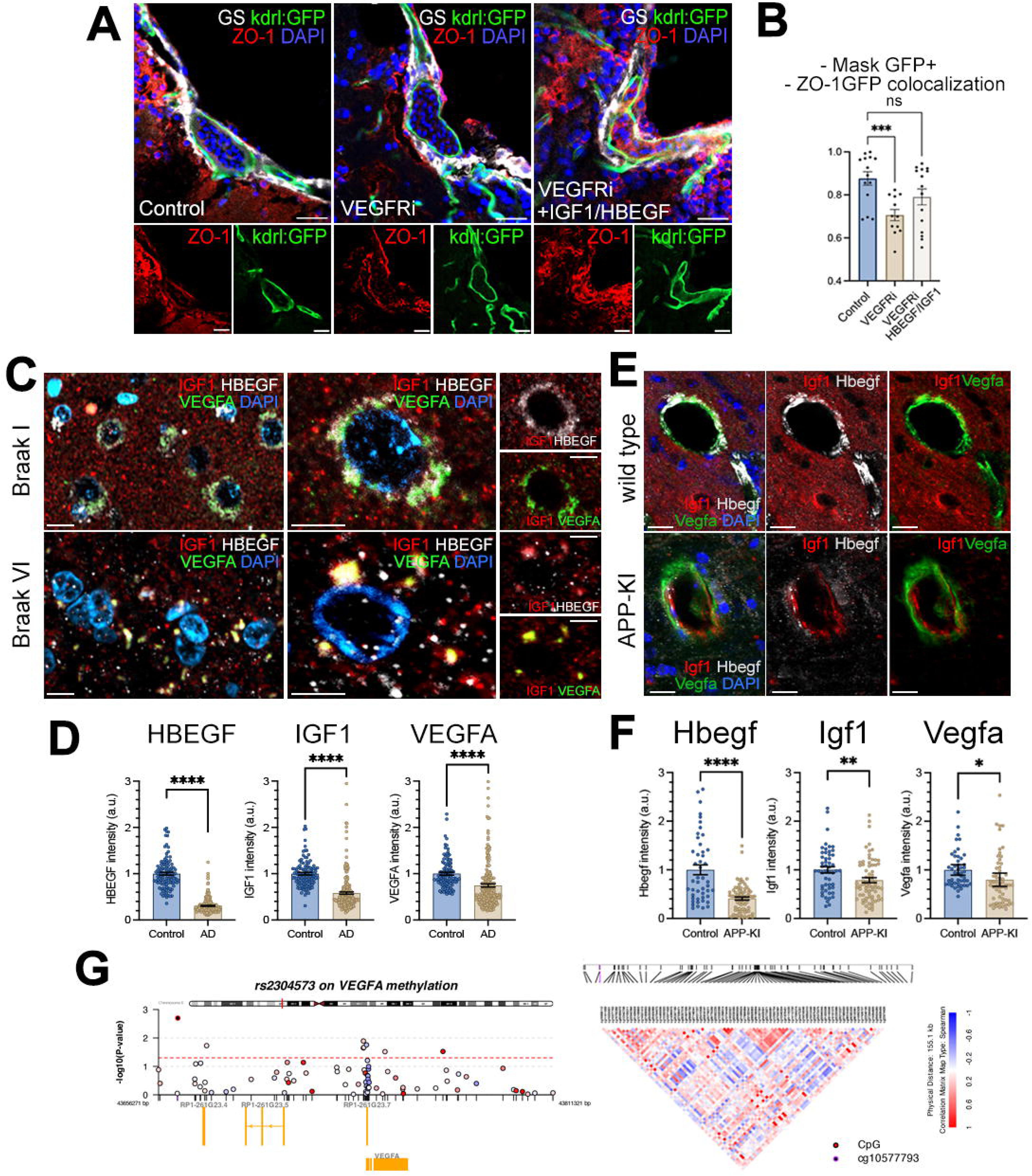
Conserved activities of HBEGF and IGF1 regulates gliovascular interactions and endothelial tight junctions. **A**, Immunostaining for GS (white), kdrl:GFP (green), ZO-1 (red) with DAPI counterstain (blue) in control, VEGFRi and VEGFRi + IGF1/HBEGF-injection in zebrafish brain. **B** Quantification graph for ZO-1/GFP colocalization, which indicates the blood-brain-barrier integrity. **C**, Immunostaining for IGF1 (red), HBEGF (white), VEGFA (green) with DAPI counterstain (blue) in postmortem control (Braak stage I) and AD (Braak stage VI) human brain. Left panels are lower magnification. Right-most panels are combinations of IGF1/HBEGF and IGF1/VEGFA. **D,** Quantification graph for HBEGF, IGF1, VEGFA intensities in human brains**. E**, IFS for IGF1 (red), HBEGF (white), VEGFA (green) with DAPI counterstain (blue) in wild type and APP knock in mouse model. **F**, Quantification graph for HBEGF, IGF1, VEGFA intensities. **G,** Regional plot showing the mQTL effects of the genetic variant of FN1 (*rs2304573*) on methylation levels of the transcriptional regulatory regions of the human VEGFA locus. Dotted red line indicates the statistical significance. Scale bars equal 25 µm. Data is represented as dot plots or multivariable graphs, and statistical significance is noted as non-significant (ns) or with the precise p-value where applicable. * (p < 0.0332), ** (p < 0.0021), *** (p < 0.002), and **** (p < 0.0001).

Gliovascular interactions, vital for BBB and brain homeostasis (*6, 7, 105, 106*), are disrupted early in AD (*2, 5, 18, 35, 107, 108*). By investigating the evolutionary conservation of VEGFA/HBEGF/IGF1 signaling axis, we observed reduced VEGFA, HBEGF, and IGF1 in zebrafish, APP-KI mice, and human AD brains (**Fig. 7C-E**). AD brains with Braak stage 5-6, which represents advanced AD pathology characterized by extensive neurofibrillary tangles in the neocortex and significant cognitive decline (*109*), also show significant alterations in VEGFA, HBEGF, and IGF1 (**Fig. 7D-F**), mirroring fibronectin-induced VEGFR/HBEGF/IGF1 signaling disruption.

To explore the molecular interplay between FN1 and VEGFA expression, we analyzed the available epigenetic datasets and methylation profiles in autopsied AD brains (*110*). We found an intronic *FN1* variant linked to altered VEGFA methylation, which supports transcriptional regulation of VEGFA by fibronectin (**Fig. 7G**). These findings align with human brain single-nucleus transcriptomics studies (*1, 2, 33, 55, 79, 80, 82, 83, 111–113*) consistently showing upregulated *FN1* gene expression and reduced VEGFA/IGF1 in astrocytes (**Supplementary Fig. 9, Supplementary Table 7**).

## Discussion

Our findings emphasize the crucial role of fibronectin as a pathological mediator in AD, with particular significance in *APOE-ε4* carriers. By integrating data from postmortem human brains, animal models, and iPSC-derived cultures, we demonstrate that fibronectin deposition contributes to BBB dysfunction through multiple interconnected mechanisms. This includes aberrant interactions with ApoE-ε4, inflammation-induced upregulation of fibronectin, and suppression of growth factor signaling pathways. These findings also highlight the therapeutic potential of a targeted approach aimed at reducing fibronectin aggregation.

Fibronectin is a key extracellular matrix protein that plays a critical role in tissue repair and remodeling (*13, 85, 114*), and is involved in the scarring process promoting the deposition of fibrous tissue and the recruitment of other matrix components to stabilize injured tissues. While this process is vital for wound healing in peripheral tissues, excessive fibronectin deposition in the central nervous system can have detrimental effects (*44, 114–117*). Fibronectin accumulation at the BBB interferes with the critical crosstalk between astrocytes and endothelia, leading to impaired clearance of metabolic waste, reduced vascular integrity, and exacerbated neuroinflammation. Such disruptions are particularly relevant in the context of AD, where the balance between repair and homeostasis is crucial to mitigating disease pathogenesis and risk. The results here mechanistically link fibronectin activity to altered gliovascular interactions and reduced BBB function.

In AD, amyloid deposition and elevated tau reduce IGF1 which mediates the effects of HBEGF expression further promoting their accumulation in brain (**Supplementary Table 10**). This suggests a mechanism that connects the combined functions of HBEGF and IGF1 and the interaction of astrocytes with endothelia to maintain a healthy BBB promoting homeostatic clearance mechanisms at the perivascular space. Interstitial clearance through the vascular basement membrane, which is primarily formed and regulated by endothelia and astrocytes, also plays a critical role in AD pathology (*118–120*). A previous study showed that HBEGF in combination with IGF1 stimulates astrocyte migration and wound closure, showcasing their synergistic impact on glial behavior during CNS injury repair (*121*). This underscores the necessity of HBEGF/IGF1 for functional astrocyte-endothelial interaction, which is critical for maintaining the structural and functional integrity of the gliovascular architecture to perform its homeostatic functions such as amyloid clearance (*5, 22, 108, 118, 120, 122*).

Our study also links FN1 activity to VEGF signaling, which is critical for maintenance of cerebrovascular health and integrity (*57, 70, 88, 123*). In AD, astrocytic states transition from maintenance roles to disease-associated reactive states, characterized by an upregulation of reactive markers and loss of homeostatic genes, including VEGFA (*83*). Previously, we showed that VEGFA, predominantly expressed by astrocytes, is significantly downregulated in AD thereby disrupting gliovascular signaling (*57, 82*). Clinically, lower VEGFA levels are associated with AD pathology such as increased tau, altered cerebral blood flow, BBB dysfunction, and impaired cognition (*57, 70, 82, 124, 125*). This aligns with the broader finding that astrocytic malfunction in AD compromise BBB integrity (*4, 6, 73*), and may augment the *APOE-ε4-*induced fibronectin activity by regulating VEGF, HBEGF and IGF1 signaling representing one of the prominent pathological mediators of AD at the BBB.

Despite the robust findings in this study, we note some limitations. Zebrafish and mouse models are useful for investigating AD and CVP, but they may not fully capture the complexity of human pathology. To further enhance the applicability of these findings to human AD, further validation in additional human brains and in reductionist models of human vascular niche may be necessary. Furthermore, the known overlap of other co-morbidities, such as diabetes and hypertension, could contribute to the observed relationships between fibronectin, VEGFA, HBEGF, IGF1, and AD pathology. Although we identified inflammation as one driver of fibronectin deposition, the specific inflammatory signals and their roles at different disease stages require further investigation.

Taken together, the observed BBB dysfunction in AD, previously linked to *APOE-ε4* (*43, 126, 127*), can now be further attributed to the pathological deposition of fibronectin, which disrupts regulatory signaling cascades by altering astrocyte-endothelial interactions, impairing the clearance of amyloid-β, and exacerbating neuroinflammation. These mechanisms underscore the multifaceted role of fibronectin in the pathogenic loss of BBB integrity because its accumulation disrupts critical regulatory cascades essential for maintaining the structural and functional integrity of BBB-associated cells. Furthermore, the *FN1* loss-of-function variant(*18*), which limits fibronectin deposition and confers protection against BBB dysfunction in *APOE-ε4* carriers, underscores the therapeutic potential of targeting fibronectin. This study establishes fibronectin as a central pathological mediator in *APOE-ε4* and AD-associated BBB dysfunction, highlighting the potential clinical benefits of preserving gliovascular integrity in mitigating AD progression.

## Materials and Methods

### Ethics statement

Human samples were sourced from the New York Brain Bank at Columbia University Vagelos College of Physicians and Surgeons and anonymized to ensure that researchers had no access to the personal information of the donors. Approval from the Institutional Review Board at Columbia University Irving Medical Center was obtained prior to the generation of clinical data. All mice used in this study were treated in accordance with the National Institutes of Health Guide for the Care and Use of Laboratory Animals and approved by the Columbia University Medical Center Institutional Animal Care and Use Committee (IACUC) (approval number AABF3553 and AABR4602). Animals were maintained under temperature and humidity-controlled environment of Columbia University Medical Center on a 12/12-h light/dark cycle, with *ad libitum* access to food and water in compliance with the protocols approved by the Institutional Animal Care and Use Committee of Columbia University (IACUC). For zebrafish, all animal experiments were performed in accordance with the applicable regulations and approved by the Institutional Animal Care and Use Committee (IACUC) at Columbia University (protocol number AC-AABN3554). Animals from the same fish clutch were randomly distributed for each experimental group. Animals were handled with caution to reduce suffering and overall animal numbers.

### Brain preparation and immunohistochemistry

Eighteen post-mortem sections of the BA9 prefrontal cortex were acquired from the New York Brain Bank at Columbia University, provided as paraffin-embedded specimens along with neuropathological evaluations. The method of Immunohistochemistry (IHC) applied followed the procedures outlined before(*20, 128*). For the IHC, primary antibodies used included FN1 (Proteintech, Cat.# 66042-1-Ig, dilution 1:250), CD31 (Abcam, Cat.# ab134168, dilution 1:250), GFAP (Thermo Fisher, Cat.# OPA1-06100, dilution 1:250), VEGF (R&D Systems, Cat.# MAB2932-100, dilution 1:200), IGF-1 (Abcam, Cat.# ab106836, dilution 1:200), HB-EGF (Thermo Fisher, Cat.# PA5119187, dilution 1:200), A-beta42 (Cell Signaling, Cat.# 8243S, dilution 1:200). Secondary antibodies employed were goat anti-mouse Alexa Fluor 448 (Thermo Fisher, Cat.# A11001, dilution 1:500),goat anti-rabbit Alexa Fluor 555 (Thermo Fisher, Cat.# A32732, dilution 1:500), donkey anti-mouse Alexa Fluor 488 (Thermo Fisher, Cat.# A32766, dilution 1:500), donkey anti-goat Alexa Fluor 555 (Thermo Fisher, Cat.# A32816, dilution 1:500), and donkey anti-rabbit Alexa Fluor 647 (Thermo Fisher, Cat.# A32795, dilution 1:500).

Human *APOE* replacement mouse models (*24, 27, 28, 30*) and APP knock in mice (APP NL-G-F (Swedish, Arctic, and Iberian) (*129*) were used for this study. Non-transgenic C57BL/6J mice served as controls for APP^NL-G-F^. APP^NL-G-F^ and control mice were given a ketamine/xylazine cocktail for deep anesthesia, and then were transcardially perfused with ice-cold 100 mM phosphate-buffered saline (PBS), pH 7.4 followed by 10% formalin (Fisher Scientific, USA). Isolated brains were incubated in 10% formalin overnight and immersed in 30% sucrose until sinking at 4°C. Brain hemispheres were cryopreserved in OCT mounting medium, sectioned into 40µm sagittal brain slices with a Leica CM3050 S cryostat, and kept in cryoprotectant at -20°C until immunofluorescence was performed. *APOE* mice were cervically dislocated, and then the brains were drop fixed for 48 hours in 10% formalin at 4 degrees. Brains were then switched to 30% sucrose at 4 degrees and cryoembedded for sectioning. 40 micron-thick coronal sections were acquired and were immunostained. Brain sections comprising of cortico-hippocampal regions from both set of experiments were selected and subjected for free-floating immunostaining. Brain sections were initially rinsed in PBS and permeabilized with 0.3% Triton X-100 (PBS-T), followed by blocking with 10% normal goat/donkey serum in PBS-T for 60 min at room temperature. Primary antibodies were applied overnight at 4°C in 2% serum in PBS-T. Following primary antibodies were used with various combinations: FN1 (Proteintech, Cat.# 66042-1-Ig, dilution 1:300), CD31 (Abcam, Cat.# ab13468, dilution 1:300), VEGF (R&D System, Cat.# MAB2932-100, dilution 1:300), IGF-1 (Abcam, Cat.# ab106836, dilution 1:300), HB-EGF (Thermo Fisher, Cat.# PA5119187, dilution 1:300). Next day, after washing with PBS-T, the sections were incubated with appropriate secondary antibodies diluted in 2% serum in PBS-T. Secondary antibodies used were goat anti-mouse Alexa Fluor 448 (Thermo Fisher, Cat.# A-11001, dilution 1:500) and goat anti-rabbit Alexa Fluor 555 (Thermo Fisher, Cat.# A-32732, dilution 1:500), donkey anti-mouse Alexa Fluor 488 (Thermo Fisher, Cat.# A-32766, dilution 1:500), donkey anti-goat Alexa Fluor 555 (Thermo Fisher, Cat.# A-32816, dilution 1:500), and donkey anti-rabbit Alexa Fluor 647 (Thermo Fisher, Cat.# A-32795, dilution 1:500). Hoechst 33342 dye (5µg/mL) (Thermo Scientific, USA) was utilized for nuclear staining for 3 min at room temperature. Following PBS-T washes, sections were mounted on Superfrost Plus slides and coverslipped using SlowFade Gold anti-fade reagent (Invitrogen, Thermo Fisher Scientific, USA). The slides were kept in the dark at 4°C until imaging. All mouse models used were at 18-22 months of age, and 3 animals from both sexes were used for every experimental group.

### Cerebroventricular microinjection (CVMI), VEGFR blockage, tissue preparation, and immunostaining in zebrafish

CVMI was performed as described (*130*). Transgenic Tg(kdrl:GFP) (*131*) zebrafish of both sexes, aged 6 months, were used for VEGFR inhibition study (**Supplementary Table 2**). All fish were sourced from the same clutch and randomly assigned to different experimental groups. The fish were exposed to a treatment regimen consisting of a solution containing Semaxanib (SU5416) at 10 µM (SelleckChem S2845), Tivozanib (AV-951) at 10 µM (SelleckChem S1207), and ZM 306416 at 10 µM (SelleckChem S2845), administered in the fish water. Doses were adjusted according to the validated dosages as described previously (*57*). This treatment was applied for 3 hours daily over a span of three consecutive days. Following the treatment, the fish were euthanized, and tissue samples were collected as described (*132–134*). Other injections include PBS (control), human amyloid-beta42 (Aβ42, 20 µM), human HB-EGF (10 μg/ml, Thermo Fisher Scientific Gibco Cat# 100-47-50UG), human IGF-1 (10 μg/ml, Thermo Fisher Scientific Gibco Cat# PHG0071), Torin2 (10 µM, Cayman Chemical Company; Cat #: 14185) and Camptothecin (1 µM, Millipore Sigma, Cat# 390238), human TNFα recombinant protein (100ng/ml, Cell Signaling Cat# 16769S), PF-573228 (100µM, FAK Inhibitor, CAS No. 869288-64-2, SelleckChem Cat# S2013). Total injection volume was 1 μl. Animals were euthanized at relevant time points and subjected either to histological tissue preparation, or to brain dissection and dissociation for single cell sequencing preparation as described (*60, 132, 135*).

For histological tissue preparation, zebrafish heads were dissected and fixed overnight at 4°C using 4% paraformaldehyde (PFA). Samples were washed several times with 0.1M phosphate buffer (PB) and incubated overnight in 20% Sucrose with 20% ethylenediaminetetraacetic acid (EDTA) solution at 4°C for cryoprotection and decalcification. The next day, samples were embedded in cryoprotectant sectioning resin OCT (Thermofisher Scientific Cat# 23-730-571) and cryosectioned into 12-μm thick sections on SuperFrost Plus glass slides. For immunohistochemistry, the sections were dried at room temperature, followed by washing steps in PBS with 0.03% Triton X-100 (PBSTx). Primary antibodies were applied overnight at 4°C. Next day, the slides were washed 3 times with PBSTx and then appropriate secondary antibodies together with DAPI were applied for 2 hours at room temperature. The slides were then washed several times before mounting with coverslips using 70% glycerol in PBS. Following primary antibodies were used with various combinations: FN1 (Proteintech, Cat.# 66042-1-Ig, dilution 1:300), GS (Abcam, Cat.# ab176562, dilution 1:500), ZO-1 (Thermo Fisher, Cat.# 33-9100, dilution 1:500), GFP (Thermo Fisher, Cat.# PA1-9533, dilution 1:2000), VEGF (R&D System, Cat.# MAB2932-100, dilution 1:500), IGF-1 (Abcam, Cat.# ab106836, dilution 1:300), HB-EGF (Thermo Fisher, Cat.# PA5119187, dilution 1:300). Secondary antibodies used were: goat anti-mouse Alexa Fluor 448 (Thermo Fisher, Cat.# A-11001, dilution 1:500), goat anti-chicken Alexa Fluor 448 (Thermo Fisher, Cat.# A-11039, dilution 1:500), goat anti-rabbit Alexa Fluor 555 (Thermo Fisher, Cat.# A-32732, dilution 1:500), goat anti-mouse Alexa Fluor 555 (Thermo Fisher, Cat.# A-21127, dilution 1:500), goat anti-rabbit Alexa Fluor 647 (Thermo Fisher, Cat.# A-32733, dilution 1:500), donkey anti-mouse Alexa Fluor 488 (Thermo Fisher, Cat.# A-32766, dilution 1:500), donkey anti-goat Alexa Fluor 555 (Thermo Fisher, Cat.# A-32816, dilution 1:500), and donkey anti-rabbit Alexa Fluor 647 (Thermo Fisher, Cat.# A-32795, dilution 1:500) For antigen retrieval of ZO-1 and FN1, slides were heated in 10 mM Sodium citrate (pH 6.0) at 85°C for 15 minutes prior to primary antibody incubation step. Imaging was performed with a Zeiss AxioImager Z1 and a Zeiss LSM800 confocal microscope. For data analysis, colocalization of immunohistochemical markers was quantified using the colocalization module in Zen software (Carl Zeiss, Germany). Statistical analysis was conducted using GraphPad Prism (GraphStats, version 6.02). Differences between experimental groups were assessed using unpaired parametric t-tests with Welch’s correction. Effect sizes were calculated using G-Power (https://www.psychologie.hhu.de/arbeitsgruppen/allgemeine-psychologie-und-arbeitspsychologie/gpower), and sample sizes were estimated with n-Query and scPower (*136*). Each experimental group included at least three animals from both genders.

### 3D hydrogel preparation, cytokine treatment, RNA isolation and sequencing

Primary Human Fetal Astrocyte (ScienCell Research Laboratories, Cat. # 1800) StarPEG-Heparin Hydrogel (3D pHA) cultures were prepared as previously described (*46, 49, 50*). Briefly, the pHAs at passage 2 were thawed and counted using an automated cell counter (Countess II FL, Thermo Fisher Scientific). 0.375 x 106 cells were plated into T75 flasks in 10 ml of complete astrocyte medium (Astrocyte medium, ScienCell Res. Lab., Cat.#. 1801, supplemented with 2% FBS, 1X Astrocyte Growth Supplement, and 1X Pen/Strep). The cells were dissociated and collected from the culture flasks using StemPro Accutase Cell Dissociation Reagent (Gibco, Cat. No. A1110501) 3 days post-thaw. After centrifugation at 271 g force (1200 rpm when using Thermo Fisher Scientific Heraeus Megafuge 16R, rotor 75003629 with 168 mm max radius) for 8 minutes, the supernatant was carefully aspirated and pHAs were resuspended in DPBS or complete AM and counted. The cell concentration was adjusted to 25,000 cells/hydrogel. Afterwards, the cells were first gently mixed with a heparin maleimide conjugate (Dr. Carsten Werner, TU Dresden, Cat.#. HM6 170530) solution and then with starPEG-MMP-peptide conjugate (Dr. Carsten Werner, TU Dresden, Cat.#. PM-CIWC-Ac 171107) solution in 1:1:2 volume ratio to obtain the desired hydrogel volume in a final concentration of 1.5 mM heparin maleimide conjugate solution and 1.12 mM starPEG-MMP-peptide conjugate solution in 20 µl final volume. Next, a 20 µl of droplet was quickly placed on a Parafilm sheet and left to gelate for about 2 min. Finally, the hydrogels were gently placed in 24-well non-tissue culture treated plates (Falcon, Cat. No. 351147) in 0.75 ml complete AM per well and cultured in a humidified cell culture incubator at 37°C. The medium was changed one day post-cell encapsulation and three times a week thereafter. The 3D pHA cultures were maintained up to 3 weeks depending on the experiment setup. Human astrocytes encapsulated in hydrogels were treated with TNFα (Peprotech, Cat.#. 300-01A, 0.1 μg/ml, 5.7471 nM), IL4 (Peprotech, Cat.#. 200-04, 0.1 μg/ml, 6.6225 nM), or a combination of both one-day post-cell encapsulation. These treatments were prepared by diluting stock solutions in complete astrocyte medium and were applied continuously by replenishing the treatment medium along with the medium of control cells three times a week until the end of three-week culture period. We used Norgen Total RNA extraction kit for isolation. In brief, 3 gels per conditions were lysed in lysis buffer followed by centrifugations and washings according to the manufacturer protocol. The concentration of RNA was measured using a NanoDrop 2000 spectrophotometer (Thermo Fisher Scientific) and quality of the RNA was assesed using an Agilent 2100 Bioanalyzer (Agilent Technologies). Samples with an RNA integrity number (RIN) greater than 9 were deemed suitable for further analysis (**Supplementary Figure 10**).

Library preparation, sequencing, aligning to the human genome and bioinformatic analyses were performed as described previously (*49, 50, 52*). In brief, cDNA libraries were prepared by following the protocol for NEBNext® Ultra I Directional RNA Library Prep Kit. This involves the following steps: mRNA isolation via poly(A)+ selection and fragmentation, first strand and second strand cDNA synthesis, purification using the Agencourt® AMPure® Kit and end repair/dA-tailing of cDNA. Adapters were ligated to the dA-tailed cDNA, followed by a size selection using AMPure XP Beads. Indexing of the library constructs was done with illumina® index primer during the following PCR amplification using NEBNext® Q5 2X PCR Master Mix. Lastly, libraries were purified using the Agencourt® AMPure® Kit. Libraries were pooled and sequenced on an illumina® NextSeq 500 system, resulting in ca. 27 – 38 million 75 bp single-end reads. All protocols are performed according to the manufacturers’ instructions. Raw sequence data were processed using a standard Illumina pipeline. Quality-controlled reads were aligned to the human genome (GRCh38) using gsnap, and differential expression analysis was conducted using DESeq2. A minimum of three biological replicates per condition was used, with power analysis ensuring adequate sample size to detect a minimum two-fold change in gene expression at a 5% error rate level.

### Cell sorting and single cell transcriptomics in zebrafish

The telencephalon of the 6- to 16-month-old fish were dissected in ice-cold PBS and directly dissociated with Neural Tissue Dissociation Kit (Miltenyi Biotec, Cat. No. 130-092-628) as described previously (*62, 132*). After dissociation, cells were filtered through 40 μM cell strainer into 10 mL 2% BSA in PBS, centrifuged at 300 g for 10 min, and resuspended in 4% BSA in PBS. Viability indicator dyes Sytox Blue (Invitrogen, Cat No. S34857) and Dycle Ruby (Invitrogen, Cat. No. V10309) were used to sort the cells by flow cytometry (FACS). Samples from different experimental groups were simultaneously sorted in separate FACS machines (Sony MA900-FP) to ensure as minimum incubation time for the samples to remain on ice. Sorting histograms are provided in **Supplementary Information 1**. The resulting single cell suspension was promptly loaded on the 10X Chromium system (*137*). 10X libraries were prepared as per the manufacturer’s instructions. Generated libraries were sequenced via Illumina NovaSeq 6000 as described (*60, 62, 132, 138, 139*). In total, 66,848 cells were sequenced for *fn1b*^-/-^ study. The raw sequencing data was processed by the Cell Ranger Single Cell Software Suite (10X Genomics, v6.1.2) with the default options. On average, 95.45% of the total 399.6 million gene reads were aligned to the zebrafish genome release GRCz11 (release 105). For single cell sequencing of the VEGFR inhibitor treatment, we treated the double transgenic (Tg(fli1a:EGFP), Tg(her4.1:RFP)) zebrafish (*86, 87*) (n=3 per group) with VEGFR inhibitor cocktail (ZM 306416 Tivozanib, Semaxanib Tivozanib, Tivozanib) or DMSO as control. Doses are adjusted according to established and validated protocol (*57*). Three days post treatment, we dissected the telencephalons and dissociated the tissue by using the same methodology as we previously described above. Filtered and dissociated cells were resuspended in 4% BSA in PBS. GFP+ and RFP+ cells were then sorted in a separate tube via Sony MA900-FP sorter, and these cells were promptly loaded on the 10X Chromium system as described above. 10X libraries were prepared, cDNA libraries were sequenced by using Cell Ranger Single Cell Software Suite (10X Genomics, v6.1.2). In total, 24,518 cells were sequenced and analyzed. On average, 95.6% of the 357.4 million gene reads per cohort mapped to the zebrafish genome release GRCz11 (release 105). The resulting matrices were used as input for downstream data analysis by Seurat (*140*).

### Read alignment, and quality control

To create Seurat (version 5) objects, we filtered out any cells with less than 200 expressed genes, and with genes expressed in less than 3 cells. After filtering out the low-quality cells, Seurat objects were normalized, and the top 2,000 variable genes were used for further analyses. Anchors were identified by using FindIntegrationAnchors function and the datasets were integrated (IntegrateData). Layers were integrated (IntegrateLayers) and joined (JoinLayers). In addition, DoubletFinder (*141*) was used to identify and remove doublets, and the rest of the analyses were done on singlets. The integrated Seurat object included 62,302 cells with 25,030 genes. The data were scaled using all genes, and 32 PCAs (RunPCA) were identified. Cell clustering, marker gene analyses, differential gene expression and preparation of feature plots were performed using Seurat V5 as described (*62, 111, 140, 142, 143*). To find differentially expressed genes, we used FindMarkers function of Seurat with 0.25 logfc.threshold. To create Seurat (version 4.1.3) objects for VEGFR inhibition study, similar analytical pipeline was used. The clusters were identified using a resolution of 1. In total, 37 clusters were identified. The main cell types were identified by using *s100b* and *gfap* for astrocytes; *kdrl* and lef1 for endothelial cells; *hbaa1* and ba1 for erythroid cells; *nell2b* and *slc17a6a* for excitatory neurons*, tubb5* and *marcksb* for immature neurons*; il2rb* and *rgs13* for immune cells*; gad1* and *gad2* for inhibitory neurons*, sv2a, nrgna, grin1a* for neurons; *cd74a* and *apoc1* for microglia; *pcna* and *mcm5* for neuroblasts; *pdgfrb* and *kcne4* for pericytes; *mbpa* and *mpz* for oligodendrocytes; *aplnra* for OPC; *myh11a* and *tagln2* for vascular smooth muscle cells (*62, 111, 135*).

### Isogenic iPSC-derived astrocytes

KOLF2.1J isogenic iPSC lines, expressing APOEε3/3 and APOEε4/4 variants, were cultured according to the described protocol (*37*). The iPSCs were maintained under feeder-free conditions on hydrogel-coated plates using mTeSR1 medium (STEMCELL Technologies). Media changes were performed daily, and cells were passaged at approximately 80% confluency to maintain optimal growth conditions. Differentiation of iPSCs into astrocytes was performed following the protocol described (*42*). Differentiation efficiency was confirmed by immunostaining for astrocytic markers such as GFAP, ensuring the generation of a homogeneous astrocytic population. For immunocytochemistry, astrocytes were fixed with 4% paraformaldehyde, permeabilized with 0.1% Triton X-100, and blocked with 5% normal serum to minimize non-specific binding. Cells were then incubated with the following primary antibodies: APOE (rabbit polyclonal antibody, Bioss, Cat #bs-5039R, 1:500), FN1 (mouse monoclonal antibody, Proteintech, Cat #66042-1-Ig, 1:500), VEGFA (mouse monoclonal antibody, R&D Systems, Cat #MAB2932, 1:500 dilution), GFAP (rabbit polyclonal antibody, Thermo Fisher, Cat #OPA1-06100, 1:500) and IGF1 (rabbit polyclonal antibody, Abcam, Cat #ab9572, 1:500). Secondary antibodies conjugated to Alexa Fluor dyes (488, 555, or 647) were used at a dilution of 1:1000, and DAPI was included for nuclear staining. iPSC-derived isogenic astrocytes were treated with human HBEGF recombinant protein (Thermo Fisher, Cat #100-47) at final concentrations of 1 ng/mL and 10 ng/mL. Treatments were performed over 24 hours, after which cells were fixed and processed for immunostaining as described above. Immunofluorescence images were acquired using a Zeiss Axio Observer microscope equipped with a confocal module. Image quantification was conducted using Zeiss Zen software, supplemented by Arivis analytical pipelines to assess marker intensity and colocalization. For quantification, images containing FN1, APOE and DAPI signals were analyzed to assess fibronectin distribution in relation to the cell nucleus, cell body and its close periphery. Masks were generated based on the DAPI+ nuclei, with boundaries extended by 10 microns from nuclear edge to encompass the cell body and its immediate periphery, capturing also the extracellular fibronectin localized near the cells. FN1 intensity within these defined regions was measured, and results were normalized to the area of the analyzed region to ensure consistency and comparability across samples. Data were then subjected to statistical analysis using GraphPad Prism. Pairwise comparisons were analyzed using t-tests with Welch’s correction, while multiple comparisons were evaluated using two-way ANOVA with Tukey’s post-hoc test. All results were expressed as mean ± SEM, and a significance threshold of p < 0.05 was applied to determine statistical relevance.

### Vascular cells, iPSC culture and cell lines

This study used APOE isogenic lines derive from 4 different donors using iPSC lines previously published (*144*). All lines were negative for mycoplasma, and karyotypically normal by G-banded Karyotype (WiCell Research Institute, Inc.). The iPSCs were maintained in six-well plates coated with growth factor-reduced Matrigel or Cultrex at 37ᵒC and 5% CO_2_. The cultures were fed with twice a week mTeSR1 medium with an FGF2DISC (Stem Cultures LLC, DSC500-48). Each week one FGF2 DISC was added into one well of a 6-well plate with 2 mL of mTeSR1 medium. The medium, but not the FGF2DISC, was replaced every 2-3 days based on culture confluency. hPSC cultures were clump-passaged using ReLeSR about once a week and a new FGF2 DISC added. Cells were not allowed to grow past 80% confluency. iPSC and vascular cells samples were genotyped periodically to confirm APOEε3/3 vs APOEε4/4 line status using Taqman SNP assays rs429358 (Life Technologies Assay ID C_3084793_20), and rs7412 (Life Technologies Assay ID C_904973_10). Assays were performed on a QuantStudio 6 Flex system.

### iPSC co-differentiation into vascular cells

When hPSCs reached approximately 80% confluency, hPSCs were passaged using Accutase and seeded at cell density of 50,000 cells/cm^2^ on Cultrex-coated tissue culture plates. hPSCs were cultured overnight in mTeSR1 medium supplemented with 10 µM Y-27632. To start the differentiation, we followed our previously published protocol (*38*). In brief, hPSC medium was replaced with LaSR medium comprised of Advanced DMEM/F12, GlutaMAX and 60 µg/mL of L-ascorbic acid and supplemented with 6 µM of CHIR99021 and cultured for 2 days to pattern cells towards mesoderm. On day 2, medium was fully replaced with LaSR medium supplemented with 100 ng/mL VEGF_165_ and 20 ng/mL FGF2.

### Cell sorting and iPSC-EC monolayer culture

Vascular differentiated cells (described above) were sorted on day 5-8 of the differentiation with CD31 magnetic beads (Miltenyi 130-091-935). Sorted hPSC-ECs were plated at densities of approximately 3.0-5.5 x10^5^ cells/cm^2^ on Embryomax (0.1% Gelatin, Sigma ES006B) coated culture plates with LaSR medium supplemented with 10% FBS, 25 ng/ml VEGF, 20 ng/ml FGF2 and 10 µM Y-27632. Medium was replaced the following day to remove Y-27632. ECs were cryopreserved in CS10 freezing medium when cultures reached about 80% confluency. For some experiments, 3 APOEε3/3 and 3 APOEε4/4 lines were thawed and cultured on embryomax coated plates for three additional days and collected for RNA. In other experiments, iPSC-ECs were not frozen down and instead cultured an additional 2-4 day post sorting with PromoCell MV2 medium (C-22022) prior to RNA isolation. In either experimental set-up, APOEε3/3 and APOEε4/4 lines were differentiated, sorted, and grown in parallel for comparisons.

### 3D iPSC vascular models in fibrin hydrogels (VAMP models)

To encapsulate unsorted vascular cells into the hydrogel, we followed methods we have established previously (*38*). In brief, vascular cells were single-cell harvested between day 5-8 of differentiation. Cells were counted, spun down and resuspended at 2×10^7^ cells/mL. Next, 25 µL of cell suspension in LaSR with thrombin (4 units/mL PBS^++^) and 25 µL of fibrinogen solution (20mg/mL in PBS^++^) were thoroughly mixed, immediately pipetted into a cell-culture treated plate. The resulting fibrin gel was comprised of 10 mg/mL of fibrinogen, 2 unit/mL of thrombin and 1×10^7^ cells/mL. After fibrin polymerization, LaSR medium was added, supplemented with 5 µL/mL of aprotinin (10,000 KIU/mL stock solution) and 50 ng/mL of VEGF to induce vessel growth. 50% medium exchanges were performed every 3-4 days. VAMPs were collected for analysis after 7 days of culture.

### 2D endothelial cell monolayers and immunofluorescence

Cells were fixed with cold 4% paraformaldehyde for 15 min and washed three times with PBS. Cells were permeabilized for 7 minutes with 0.1% Saponin (Sigma S7900) in PBS for 7 min, washed with PBS and blocked in 2% Fish gelatin for 45 min. Antibody buffer was 0.01% Saponin in 0.5% Fish gelatin in PBS. Primary antibodies done in series with wash between (CD31 BD Biosciences 550389, dilution 1:200) and ZO1 (Protein tech cat# 21773-1-AP, lot 00106957, dilution factor 1:200) are 12 hours at 4C or 2 hr at RT on rocker. Washes between steps are at RT on rocker with PBS for 5 min, 2 times, followed by one wash with antibody buffer. Secondary antibodies (1:250, AF568 anti-mouse A11019, AF647 anti-rabbit Jackson cat# 705-606-147) are 45 min at RT on rocker shaker, done in series, followed by DAPI for 15 min. Fluorescence imaging is completed on Echo Revolution microscope with 20x objective (NA 0.45), equipped with DAPI, FITC, Texas Red and Cy5 em/ex filters. Images were converted to OME.Tiff. Images shown are cropped at higher zoom for visualization.

### 3D VAMP whole mount immunofluorescence

Cells were fixed with cold 4% paraformaldehyde for 1 hour on a rocker at 4C and washed three times with PBS. Antibody blocking buffer was comprised of 10% Normal Goat Serum, and 3% Bovine Serum Albumin in dPBS. Primary antibodies (CD31, Fisher BDB550389, 1:200; FN1, Thermo PA5-29578, dilution factor) were incubated overnight, washed 3 times with PBS over 8 hours and secondary incubated overtime followed by another 3 washes. Primary antibody stains were done in series (i.e., CD31 and then FN1 staining). 5 images per model were taken with an Echo Revolution microscope using a 10x/0.3 Fluorite objective. Pseudo Z sections were taken every 5.192 microns. Z-stacks were deconvolved using ImagePro11 software and projected and processed using ImarisViewer.

### RNA isolation, cDNA preparation, and qPCR in iPSC-ECs

Total RNA was isolated from cells with Zymo RNA Clean and Concentrator Kit. RNA concentrations were measured on a nanodrop, and cDNA was generated using High-Capacity cDNA Reverse Transcription Kit. qPCR was run on 1.25ng cDNA/reaction using Taqman probes for FN1 (Assay ID Hs01549976_m1, Life Technologies) with 18S (Assay ID Hs01026310_m1, Life Technologies) as a loading control. qPCR was run on a QuantStudio 6 Flex system.

### Quantitative real-time PCR in zebrafish

HBEGF was microinjected in the adult telencephalon of reporter transgenic zebrafish line Tg(her4:GFP) (*145*) that labels GFP-positive astrocytes. One day after cerebroventricular microinjection, the brains were dissected, and single cell suspensions were generated as described above. Following the FACS, GFP-positive sorted astrocytes and GFP-negative sorted remaining cells mainly neurons were immersed in TRIzol™ Reagent (Invitrogen, Cat.No: 15596026). Sorting histograms are provided in **Supplementary Information 1.** RNA isolation was performed by using Direct-zol RNA Microprep Kit (Zymo Research, Cat.No: R2060) according to the manufacturer’s instructions. Complementary DNA (cDNA) was obtained from the purified RNA using the SuperScript™ III First-Strand Synthesis System (cat. no. 18080051; Invitrogen; Thermo Fisher Scientific, Inc.) with oligo (dT) by following the manufacturer’s protocol. The RT-qPCR reaction was performed using a final volume of 10 μl. To reach the final volume, we used 5 μl of PowerUp™ SYBR™ Green Master Mix 2x (Applied Biosystems, Cat.No.: A25742), 1 μM of each primer (*igf1* forward #1: CAGCAAACCGACAGGATATGG, reverse #1: CAGCTCTGAAAGCAGCATTCG from ref.(*146*); igf1 forward #2 GGGATGTCTAGCGGTCATTT, reverse #2: CAGTGAGAGGGTGTGGGTA; *vegfa* forward: TTCGAGCGCCTCATCATTAC, *vegfa* reverse: GCTGCTGGTAGACATCATCC), beta-actin forward: ATGCAGAAGGAGATCACATCCC, beta-actin reverse: GCTTGCTAATCCACATCTGCTG, 1 μl of the cDNA (10 ng in final concentration) and UltraPure™ DNase/RNase-Free Distilled Water dH2O (Invitrogen™ Cat.No.: 10977015). The cycle threshold (Ct) value for the target gene was calculated using QuantStudio^TM^ 3 real-time PCR system. Amplification conditions for RT-qPCR were as follows: 50 °C for 2 mins, 95 °C for 10 mins, followed by 40 cycles at 95 °C (15 secs) and 60 °C (1 min). Melting curve was analyzed for each sample by following 95 °C for 15 secs, 60 °C for 1 min and 95 °C for 1 sec after amplification reactions.

### Human single nucleus datasets

Pre-processing and quality control steps for snRNA-seq data by library alignment and background noise removal, de-multiplexing, normalization and clustering, removal of low-quality cells, and cell type classification was performed as described (*54, 57*). We tested the association between pathological AD and normalized snRNA-seq data by logistic regression, while controlling for age and sex. Results were corrected for multiple hypothesis testing by calculating the false discovery rate (FDR).

### Human brain DNA methylation measurement

The genome-wide DNA methylation profile was measured by the Infinium MethylationEPIC Kit (Illumina) on New York Brain Bank, NIA AD-FBS/NCRAD and EFIGA cohorts (*147*). On the sample level quality control (QC), we have checked the control probes, sex mismatch, contamination, and genotype outlier calling to identify and remove those samples failed any of these QC metrics. On the CpG probe level QC, we kept those CpG sites with detection *P* value < 0.01 across all the qualified samples and mask those sample specific CpG site with new detection *P* > 0.01 (*148*). We further removed those CpG sites reported to have cross-hybridization problems (*149, 150*) and those polymorphic CpG sites (*150, 151*). We further corrected the dye bias for all the qualified CpG probes. For this study, we used 20 Hispanic samples with both DNA methylation and Genome-wide association study (GWAS) data.

### Causal mediation analyses

Causal mediation modeling in ROSMAP cohort RNA sequencing samples (*152*) to test whether gene expression of mediator mediates the association between gene expression of predictor and trait in ROSMAP DLPFC samples was conducted utilizing the mediation package in R, and confidence intervals were established through a nonparametric bootstrap method involving 1000 resamples (*153*).

### Image acquisition, quantification, statistical analyses

For each histological section, five randomly chosen field images were captured from immunostained slides utilizing the Zeiss LSM800 confocal microscope, which features ZEN software (blue edition, version 3.2, Carl Zeiss, Jena, Germany). Vascular markers were used to outline blood vessels in coronal sections via ZEN software’s selection tool, from which measurements of fluorescence intensity, diameter, and area were derived. The image capturing process was conducted blindly, with identifiers, neuropathological information, and genotypic data disclosed only post-acquisition, and vessels were excluded only if their diameters exceeded 50 μm. Statistical analysis was conducted using GraphPad Prism software, version 9.2.0. One-way Brown-Forsythe and Welch ANOVA tests, employing a two-stage linear step-up procedure by Benjamini, Krieger, and Yekutieli for multiple comparisons and individual variance calculations, were utilized. Kruskal-Wallis test with Dunn’s multiple comparison test was applied for non-Gaussian data sets. Unpaired two-tailed parametric t-test with Welch’s correction was used for pairwise comparisons for datasets with unequal standard deviations. For mouse brain sections, five randomly chosen field images from at least three histological sections were captured from each animal. Analysis for mouse images was similar to that of human brain analysis as mentioned above and Kolmogorov-Smirnov test was applied for statistical analysis of mouse datasets. For zebrafish brain sections, at least 5 images were acquired from at least three animals per experimental conditions. For fluorescence intensity colocalization, Manders colocalization coefficient (*154*) was calculated using ZEN software colocalization module (blue edition, version 3.2, Carl Zeiss, Jena, Germany). All statistical significances in image quantifications were denoted by * (p < 0.0332), ** (p < 0.0021), *** (p < 0.002), and **** (p < 0.0001), with the absence of asterisks indicating a lack of significance (ns). No sample sets were omitted from analysis, except in cases where histological sections were significantly damaged during preparation.

## Data availability statement

All sequencing datasets generated in this study are deposited to NCBI Gene expression Omnibus (GEO) with the following accession numbers: GSE268721, GSE268780, GSE268803, and GSE269111. Previous datasets used in this study are GSE108038 and GSE78117.

## Author contributions

PB, EY, BNV, RM, and CK conceived and designed this study. PB, EY, and CK performed the human and zebrafish brain immunolabeling, quantitative assays, image acquisition, quantification, and statistical analyses. EY, PB, CK analyzed the single cell transcriptomics in zebrafish. NN performed zebrafish brain tissue sectioning. AFT provided the human brain samples. AJL, BV, RM provided human datasets and analytical samples. YM performed methylation analyses. VCH and PDJ provided microglial activity blocking compound. XW, OI, NET integrated human single nucleus datasets. WL and YZ provided the TNFα blocking compound. TN provided mouse brain tissue. HYK and PB prepared the mouse brain tissues and performed staining. SH, DJ provided the *fn1b* knockout zebrafish. HC conducted 3D pHA experiments and analyses. HC, MIC, CK performed bulk RNA sequencing in 3D cultures and data analyses. EOC performed isogenic iPSC-derived astrocyte cultures, astrocytic differentiation, immunocytochemistry, imaging, and quantification. TB, ESF, KT and ST performed 2D iPSC-EC and 3D VAMP modeling of isogenic line pairs and collection of relevant data. All authors contributed to the interpretation of the results. PB, EY, BNV, RM, and CK generated the figures and wrote the original manuscript. All authors contributed to the editing of the manuscript.

## Acknowledgements

We would like to thank Molecular Pathology (MPSR) and Flow Cytometry Core Facility (CCTI, supported in part by the Office of the Director, National Institutes of Health under awards S10OD020056) platforms of the Columbia University Herbert Irving Comprehensive Cancer Center for procedural support, New York Brain Bank for post-mortem human brain sections, Taub Institute Imaging Platform, and DRESDEN-Concept Genome Center for total RNA-sequencing of 3D cultures or pHAs. We thank the contributors, who collected samples used in this study; the patients and families for their participation, without whom these studies would not have been possible. We thank Drs. Syed Abid Hussaini and Gustavo A. Rodriguez for providing brain tissue from aged APP-KI mice, Drs. Bengisu Turgutalp and Sherida de Leeuw for initial help with the optimization of the isogenic iPSC cultures, Drs. Uwe Freudenberg and Carsten Werner for starPEG-Heparin hydrogel materials, Delaney Flaherty (New York Brain Bank) for postmortem brain samples, Michael Kissner (Columbia Stem Cell Initiative Flow Cytometry Core Facility) for his help during sorting, and Erin Bush (Single Cell Analysis Core and Columbia Genome Center, Sulzberger Genome Center).

This work was supported by National Institute on Aging R01 AG067501 (Genetic Epidemiology and Multi-Omics Analyses in Familial and Sporadic Alzheimer’s Disease Among Secular Caribbean Hispanics and Religious Order) (RM, BNV, CK) and National Institute on Aging RF1 AG066107 Epidemiological Integration of Genetic Variants and Metabolomics Profiles in Washington Heights Columbia Aging Project (RM, BNV, CK), Schaefer Research Scholars Award (CK), Taub Institute Grants for Emerging Research (TIGER) (CK), Thompson Family Foundation Program for Accelerated Medicine Exploration in Alzheimer’s Disease and Related Disorders of the Nervous System (TAME-AD) (CK), Carol and Gene Ludwig Family Foundation (CK, AJL, BNV), Toffler Scholar Program (PB), and P30 AG066462 Alzheimer’s Disease Research Center (AFT), The content of this publication is solely the responsibility of the authors and does not necessarily represent the official views of the National Institutes of Health.

The National Institute on Aging-AD Family-based study (NIA AD-FBS; https://www.neurology.columbia.edu/research/research-centers-and-programs/national-institute-aging-alzheimers-disease-family-based-study-nia-ad-fbs) collected the samples used in this study and is supported by National Institute on Aging (NIA) grants U24AG026395, U24AG021886, R01AG041797, and U24AG056270. Additional families were contributed to the NIA-AD FBS through NIH grants: R01AG028786, R01AG027944, RO1AG027944, RF1AG054074, U01AG052410. The NIA-AD FBS began in 2003 with the goal of recruiting large, multiply affected families with late-onset Alzheimer’s disease (AD) for genetic research. The study created a resource of well-characterized families with late-onset AD. The initial phases of the Alzheimer’s Disease Sequencing Project (ADSP) included genotyping of hundreds of participants from NIA-AD FBS. The ADSP Follow-Up Study heavily engages resources provided by the NIA-AD FBS and depends upon the longitudinal follow-up of families, and the collection of additional families, in particular those from diverse populations. Samples include biological materials for genome wide association studies (GWAS) and whole genome sequencing (WGS), peripheral blood mononuclear cells (PBMC) for stem cell modeling, plasma for studies of metabolomics, proteomics, and biomarker research, and brain autopsy materials for bulk RNA sequencing.

Estudio Familiar de Influencia Genetica en Alzheimer (EFIGA) is a study of sporadic and familial Alzheimer’s Disease among Caribbean Hispanics recruited from clinics in the Dominican Republic and New York (R01 AG067501). The goal of this study is to identify genetic variants that increase late onset Alzheimer disease risk in this ethnic group. This study was initiated in 1998 and recruited individuals and their families in New York as well as from clinics in the Dominican Republic. Recruitment for the EFIGA began in 1998, to study the genetic architecture of AD in the Caribbean Hispanic population. Patients with familial AD were recruited and if a sibling of the proband had dementia, all other living siblings and available relatives underwent evaluation. Cases were defined as any individual meeting NINCDS-ADRDA criteria for probable or possible AD.

The results published here are in part based on data obtained from the AD Knowledge Portal (https://adknowledgeportal.org). The Mayo RNAseq study data was led by Dr. Nilüfer Ertekin-Taner, Mayo Clinic, Jacksonville, FL as part of the multi-PI U01 AG046139 (MPIs Golde, Ertekin-Taner, Younkin, Price) using samples from The Mayo Clinic Brain Bank.

## Supplementary Information

**Supplementary Fig. 1: Bulk RNA sequencing in 3D hydrogel cultures. A,** Sample correlation graph for control, IL4-treated, TNFα-treated, and TNFα+IL4-treated replicates. **B**, Principal component analyses of groups. **C**, MA plots for comparison between samples. Red, overexpression above significance threshold, green, downregulated above significance threshold.

**Supplementary Fig. 2: TNFα treatment response. A,** KEGG pathway schematics indicating changes NFkB and TNFα signaling components after TNFα treatment in 3D primary human astrocytes (pHA) cultures. Red: upregulated, green: downregulated. **B**, KEGG pathway chart for differentially expressed genes in TNFα-treated versus control 3D pHA cultures and TNFα+IL4-treated versus TNFα-treated 3D pHA cultures. **C**, Immunostaining for GFAP with DAPI counterstain in control and TNFα-treated 3D pHA cultures. Scale bars equal 25 µm.

**Supplementary Fig. 3: TNFα and IL4 receptor expression in 3D pHA cultures.** Sequencing read numbers of *TNFRSF1A, TNFRSF1B* and *IL4R* in 3D pHA cultures in control and after TNFα-treatment.

**Supplementary Fig. 4: TNFα activates NFkB signaling in zebrafish brain. A,** In vivo experimental scheme. Adult transgenic reporter zebrafish that express GFP under the NFkB promoter were injected. Histological sections were prepared at 1 day after the injection. Immunostaining for GFP is followed by quantification and statistical analyses of GFP-positive cells in the medial telencephalic region. **B-E**, GFP and DAPI staining on brain sections of NFkB:GFP reporter transgenic animals injected with PBS (**B, C**) and TNFα (**D, E**). **B’,** Individual GFP fluorescence channel of B. **D’,** Individual GFP fluorescence channel of D. **C, E,** Topological fluorescence histograms for NFkB activity in the brain sections of the animals injected with PBS (**C**) and TNFα (**E**). **F**, Quantification graph for average number of NFkB-positive cells per telencephalic sections under different treatment conditions in **B-D**. **G-K,** GFP and DAPI staining on brain sections of NFkB:GFP reporter transgenic animals injected with PBS (**G**), TNFα (**I**), and TNFα + T17 (TNFα blocker, ref. (*155*)) **H, J, L**, Topological fluorescence histogram for NFkB activity in **G** (**H**), **I** (**J**), and **L** (**K**). **M**, Quantification graph for average number of NFkB-positive cells per telencephalic sections under different treatment conditions in **G-K**. n = 3 animals for every experimental group, and 15 brain sections in total. Statistical analyses for multiple comparisons were performed with Dunnett’s (**F**) and Bonferroni’s (**M**) multiple comparison tests. **: p<0.01, ***: p<0.005. Scale bars equal 50 µm.

**Supplementary Fig. 5: Single cell transcriptomics in *fn1b* homozygous knockout animals after Aβ42. A,** UMAP for 38 clusters. **B**, Dot plot for cell type markers. **C**, UMAP with cell types color-coded. **D**, Violin plots for genes per cell, reads per cell, percent mitochondrial gene expression and percent ribosomal gene expression. **E**, Distribution graphs for percent mitochondrial gene expression against reads per cell in wt + Aβ42 and *fn1b*^-/-^ + Aβ42 samples. **F,** Distribution graphs for number of sequenced genes per cell against reads per cell in wt + Aβ42 and *fn1b*^-/-^ + Aβ42 samples. **G**, Distribution plot for most variable gene expressions. **H**, Total number of cells sequenced and their pie chart distribution to cell types. **I**, Heat map of comparing percentages of sequenced cell types in wt + Aβ42 and *fn1b*^-/-^ + Aβ42 samples.

**Supplementary Fig. 6: Expression of VEGF ligands. A,** Violin plots for expression of *vegfa, vegfb, vegfc*, and *vegfd* in zebrafish brain. **B,** Split violin plots for expression of *vegfa, vegfb, vegfc*, and *vegfd* comparing control and Aβ42 treatment conditions. Data from refs.(*18, 60, 62, 135*).

**Supplementary Fig. 7: Expression of integrin receptors in 3D pHA cultures and adult zebrafish brain astrocytes.** Heat maps for bulk RNA sequencing reads in 3D pHA and her4:GFP-positive sorted astrocytes from zebrafish brain. Data from refs.(*50, 60, 62*).

**Supplementary Fig. 8: Single cell transcriptomics in VEGFRi zebrafish brains. A,** Dot plot for cell type markers. **B**, UMAP for cell types in color-codes. **c**, UMAP for cell types in control (left) and VEGFRi (right) samples. **D**, Total number of cells sequenced per experimental group as pie charts, and their cell type percentages. **E**, Heat map comparing sequenced cell numbers in clusters (control and treatment - VEGFRi). **F**, Distribution graphs for percent mitochondrial gene expression against reads per cell in control and VEGFRi samples. **G,** Distribution graphs for number of sequenced genes per cell against reads per cell in control and VEGFRi samples. **H**, Distribution plot for most variable gene expressions. **I**, Violin plots for sequenced genes per cell, reads per cell, percent mitochondrial gene expression and percent ribosomal gene expression. Vehicle-treated and VEGFRi fli1a:EGFP-sorted cells had two replicates, vehicle-treated and VEGFRi her4:RFP-sorted cells has one replicate.

**Supplementary Fig. 9: Expression of FN1, VEGFA, IGF1 in integrated human single nucleus dataset**. Graph indicating astrocyte clusters in human single nucleus datasets that upregulate *FN1* and downregulate *IGF1* and *VEGFA*. Data derived from integration of single nucleus datasets in refs. (*2, 33, 57, 79*) by ref. I: ref.(*57*) Y: ref.(*2*)., Z: ref. (*33*), M: ref. (*79*).

**Supplementary Fig. 10: RNA quality in bulk RNA sequencing of 3D pHA cultures.** Electrophoresis file run summary and RNA integrity measures by electropherogram summaries and RIN values.

**Supplementary Table 1:** Source data.

**Supplementary Table 2:** Differential expression dataset for 3D hydrogel cultures after TNFα treatment.

**Supplementary Table 3:** Differential expression dataset for 3D hydrogel cultures after IL4 treatment.

**Supplementary Table 4:** Differential expression dataset for 3D hydrogel cultures after TNFα + IL4 treatment.

**Supplementary Table 5:** Differential expression dataset in control and *fn1b* knockout zebrafish brains after Aβ42.

**Supplementary Table 6:** Differential expression dataset in control and VEGFR inhibitor treated (VEGFRi) zebrafish brains.

**Supplementary Table 7:** *VEGFA* expression in integrated human single nucleus datasets comparing AD to control brains.

**Supplementary Table 8:** Association of FN1 variation *rs13004592* to changes in methylation sites of VEGFA locus in humans.

**Supplementary Table 9**: Causal mediation analyses between *HBEGF* and *IGF1* for AD traits in ROSMAP brain samples.

**Supplementary Information 1**: Flow cytometry gating parameters.

## Notes

### Competing Interest Statement

The authors have declared no competing interest.

