## Supplementary Figures for "*APOE-*ε4-induced Fibronectin at the blood-brain barrier is a conserved pathological mediator of disrupted astrocyte-endothelia interaction in Alzheimer’s disease"

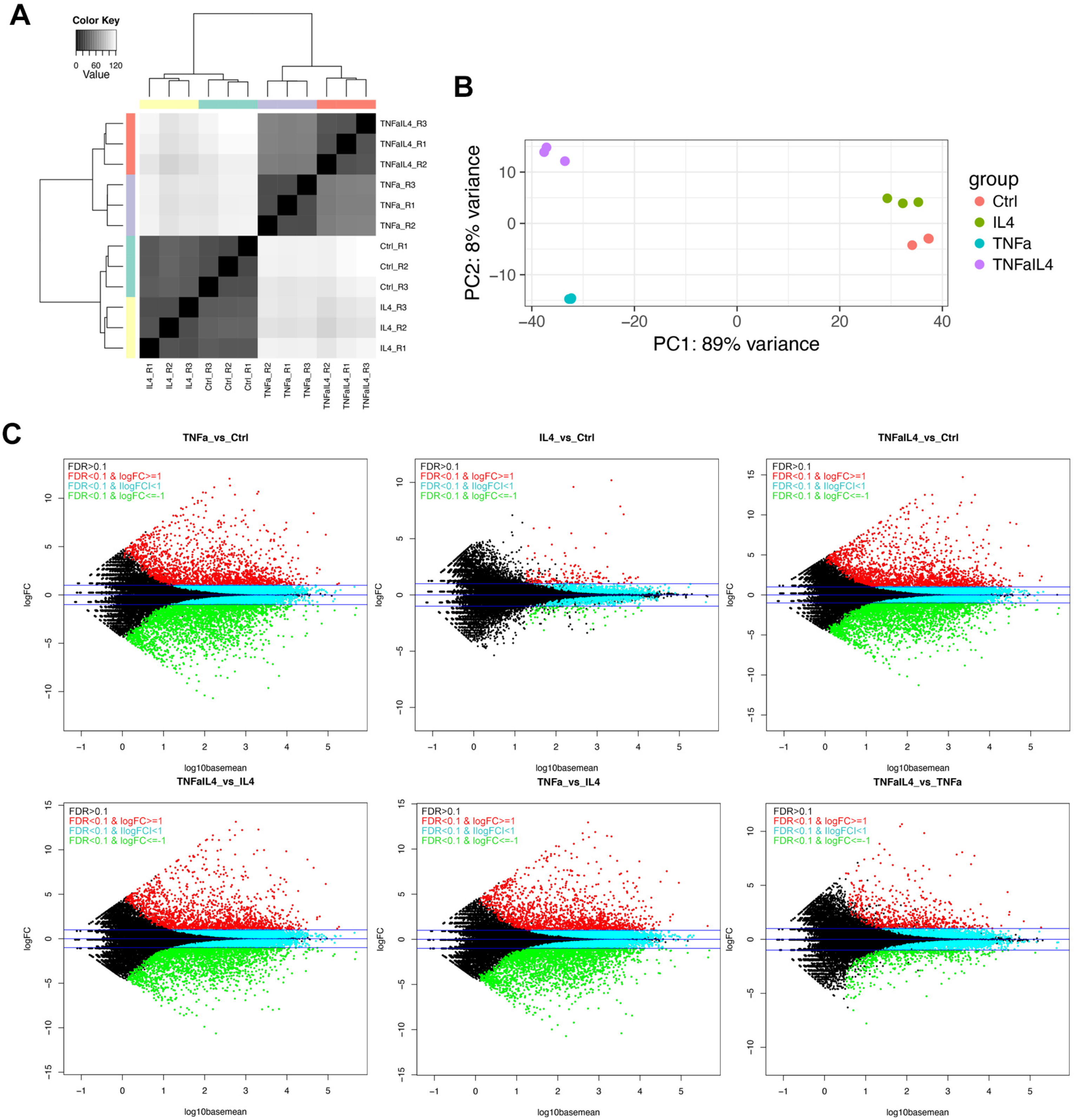

A
Color Key
0 60 120
Value
TNFaIL4_R3
TNFaIL4_R1
TNFaIL4_R2
TNFa_R3
TNFa_R1
TNFa_R2
Ctrl_R1
Ctrl_R2
Ctrl_R3
IL4_R3
IL4_R2
IL4_R1
IL4_R1
IL4_R2
IL4_R3
Ctrl_R3
Ctrl_R2
Ctrl_R1
TNFa_R2
TNFa_R1
TNFa_R3
TNFaIL4_R2
TNFaIL4_R1
TNFaIL4_R3
B
PC2: 8% variance
PC1: 89% variance
group
Ctrl
IL4
TNFa
TNFaIL4
C
TNFa_vs_Ctrl
IL4_vs_Ctrl
TNFaIL4_vs_Ctrl
TNFaIL4_vs_IL4
TNFa_vs_IL4
TNFaIL4_vs_TNFa
FDR>0.1
FDR<0.1 & logFC>=1
FDR<0.1 & llogFCl<1
FDR<0.1 & logFC<=-1
logFC
log10basemean

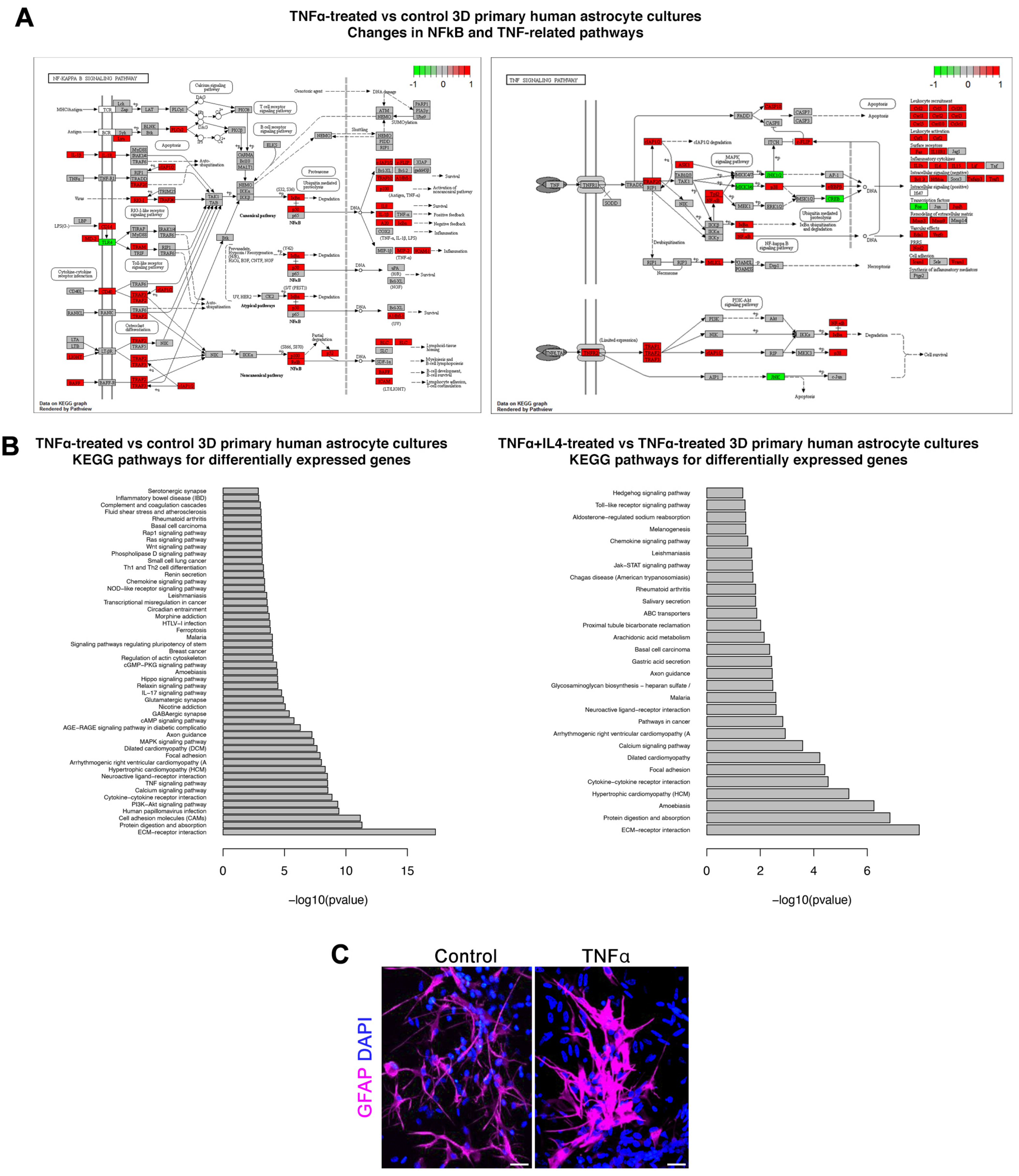
A
TNFα-treated vs control 3D primary human astrocyte cultures
Changes in NFkB and TNF-related pathways
NF-KAPPA B SIGNALING PATHWAY
TNF SIGNALING PATHWAY
-1 0 1
Data on KEGG graph
Rendered by Pathview
B
TNFα-treated vs control 3D primary human astrocyte cultures
KEGG pathways for differentially expressed genes
Serotonergic synapse
Inflammatory bowel disease (IBD)
Complement and coagulation cascades
Fluid shear stress and atherosclerosis
Rheumatoid arthritis
Basal cell carcinoma
Rap1 signaling pathway
Ras signaling pathway
Wnt signaling pathway
Phospholipase D signaling pathway
Small cell lung cancer
Th1 and Th2 cell differentiation
Renin secretion
Chemokine signaling pathway
NOD−like receptor signaling pathway
Leishmaniasis
Transcriptional misregulation in cancer
Circadian entrainment
Morphine addiction
HTLV−I infection
Ferroptosis
Malaria
Signaling pathways regulating pluripotency of stem
Breast cancer
Regulation of actin cytoskeleton
cGMP−PKG signaling pathway
Amoebiasis
Hippo signaling pathway
Relaxin signaling pathway
IL−17 signaling pathway
Glutamatergic synapse
Nicotine addiction
GABAergic synapse
cAMP signaling pathway
AGE−RAGE signaling pathway in diabetic complicatio
Axon guidance
MAPK signaling pathway
Dilated cardiomyopathy (DCM)
Focal adhesion
Arrhythmogenic right ventricular cardiomyopathy (A
Hypertrophic cardiomyopathy (HCM)
Neuroactive ligand−receptor interaction
TNF signaling pathway
Calcium signaling pathway
Cytokine−cytokine receptor interaction
PI3K−Akt signaling pathway
Human papillomavirus infection
Cell adhesion molecules (CAMs)
Protein digestion and absorption
ECM−receptor interaction
0 5 10 15
−log10(pvalue)
TNFα+IL4-treated vs TNFα-treated 3D primary human astrocyte cultures
KEGG pathways for differentially expressed genes
Hedgehog signaling pathway
Toll−like receptor signaling pathway
Aldosterone−regulated sodium reabsorption
Melanogenesis
Chemokine signaling pathway
Leishmaniasis
Jak−STAT signaling pathway
Chagas disease (American trypanosomiasis)
Rheumatoid arthritis
Salivary secretion
ABC transporters
Proximal tubule bicarbonate reclamation
Arachidonic acid metabolism
Basal cell carcinoma
Gastric acid secretion
Axon guidance
Glycosaminoglycan biosynthesis − heparan sulfate /
Malaria
Neuroactive ligand−receptor interaction
Pathways in cancer
Arrhythmogenic right ventricular cardiomyopathy (A
Calcium signaling pathway
Dilated cardiomyopathy
Focal adhesion
Cytokine−cytokine receptor interaction
Hypertrophic cardiomyopathy (HCM)
Amoebiasis
Protein digestion and absorption
ECM−receptor interaction
0 2 4 6
−log10(pvalue)
C
Control
TNFα
GFAP DAPI

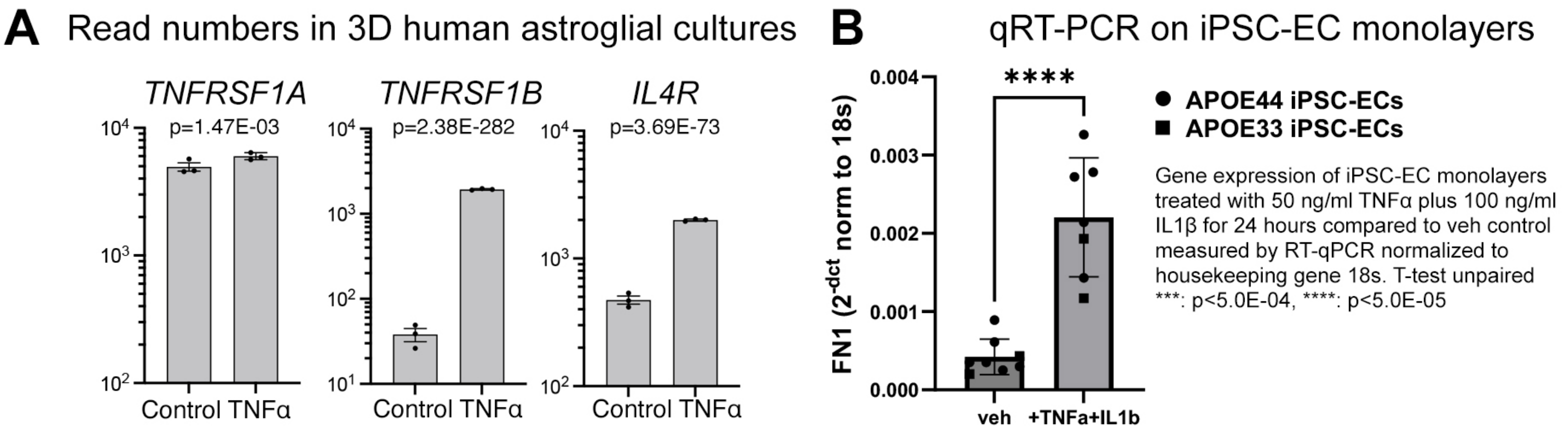
A
Read numbers in 3D human astroglial cultures
TNFRSF1A
p=1.47E-03
TNFRSF1B
p=2.38E-282
IL4R
p=3.69E-73
Control TNFα
B
qRT-PCR on iPSC-EC monolayers
FN1 (2-dct norm to 18s)
****
veh
+TNFa+IL1b
APOE44 iPSC-ECs
APOE33 iPSC-ECs
Gene expression of iPSC-EC monolayers treated with 50 ng/ml TNFα plus 100 ng/ml IL1β for 24 hours compared to veh control measured by RT-qPCR normalized to housekeeping gene 18s. T-test unpaired ***: p<5.0E-04, ****: p<5.0E-05

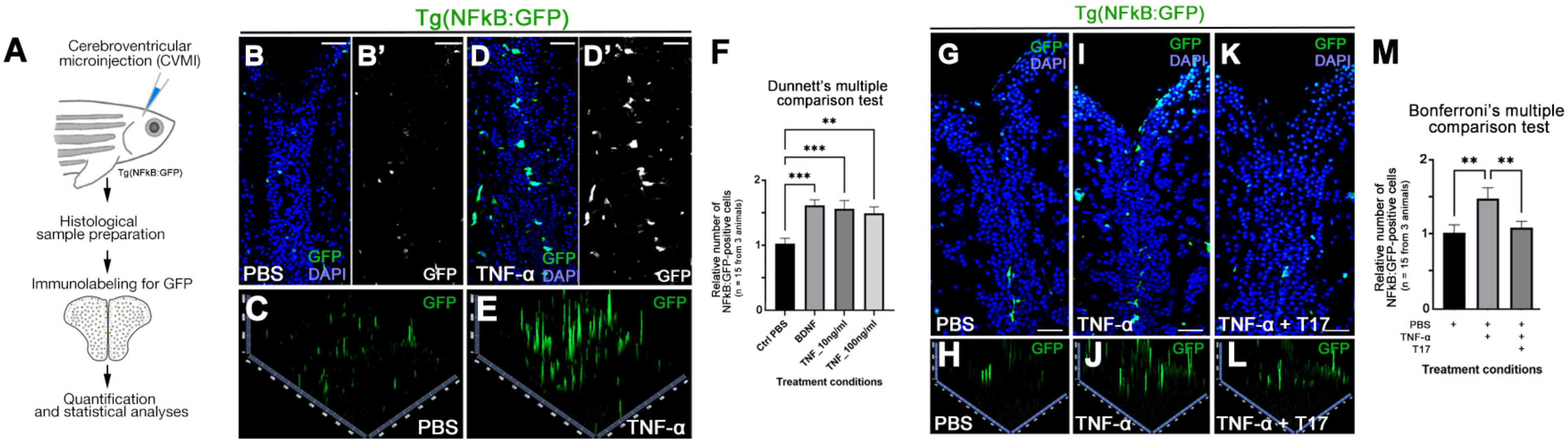
A
Cerebroventricular
microinjection (CVMI)
Tg(NFkB:GFP)
Histological
sample preparation
Immunolabeling for GFP
Quantification
and statistical analyses
Tg(NFkB:GFP)
B
B'
D
D'
GFP
DAPI
PBS
GFP
TNF-α
GFP
DAPI
GFP
C
GFP
PBS
E
GFP
TNF-α
F
Dunnett's multiple
comparison test
Relative number of
NFkB:GFP-positive cells
(n = 15 from 3 animals)
Ctrl PBS
BDNF
TNF_10ng/ml
TNF_100ng/ml
Treatment conditions
Tg(NFkB:GFP)
G
I
K
GFP
DAPI
PBS
TNF-α
TNF-α + T17
H
J
L
GFP
PBS
TNF-α
TNF-α + T17
M
Bonferroni's multiple
comparison test
Relative number of
NFkB:GFP-positive cells
(n = 15 from 3 animals)
PBS
TNF-α
T17
Treatment conditions

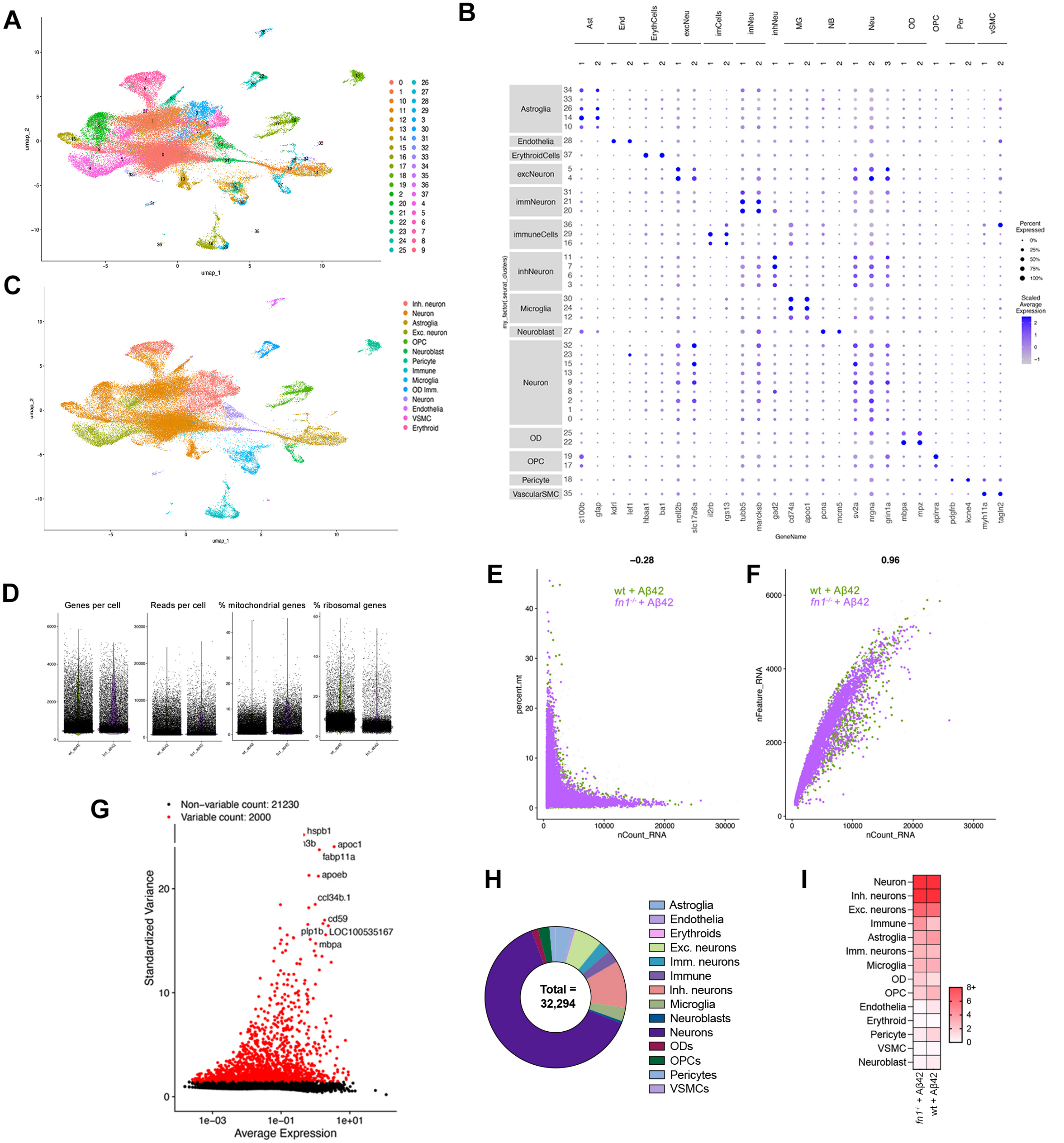
A
umap_1
umap_2
B
Ast
End
ErythCells
excNeu
imCells
imNeu
inhNeu
MG
NB
Neu
OD
OPC
Per
vSMC
Astroglia
Endothelia
ErythroidCells
excNeuron
immNeuron
immuneCells
inhNeuron
Microglia
Neuroblast
Neuron
OD
OPC
Pericyte
VascularSMC
my_factor(seurat_clusters)
s100b
gfap
kdrl
lef1
hbaa1
ba1
nell2b
slc17a6a
il2rb
rgs13
tubb5
marcksb
gad2
cd74a
apoc1
pcna
mcm5
sv2a
nrgna
grin1a
mbpa
mpz
aplnra
pdgfrb
kcne4
myh11a
tagln2
GeneName
Percent Expressed
0%
25%
50%
75%
100%
Scaled Average Expression
C
Inh. neuron
Neuron
Astroglia
Exc. neuron
OPC
Neuroblast
Pericyte
Immune
Microglia
OD Imm.
Neuron
Endothelia
VSMC
Erythroid
umap_1
umap_2
D
Genes per cell
Reads per cell
% mitochondrial genes
% ribosomal genes
E
−0.28
wt + Aβ42
fn1-/- + Aβ42
percent.mt
nCount_RNA
F
0.96
wt + Aβ42
fn1-/- + Aβ42
nFeature_RNA
nCount_RNA
G
Non−variable count: 21230
Variable count: 2000
hspb1
apoc1
fabp11a
apoeb
ccl34b.1
cd59
plp1b
LOC100535167
mbpa
Standardized Variance
Average Expression
H
Total = 32,294
Astroglia
Endothelia
Erythroids
Exc. neurons
Imm. neurons
Immune
Inh. neurons
Microglia
Neuroblasts
Neurons
ODs
OPCs
Pericytes
VSMCs
I
Neuron
Inh. neurons
Exc. neurons
Immune
Astroglia
Imm. neurons
Microglia
OD
OPC
Endothelia
Erythroid
Pericyte
VSMC
Neuroblast
8+
6
4
2
0
fn1-/- + Aβ42
wt + Aβ42

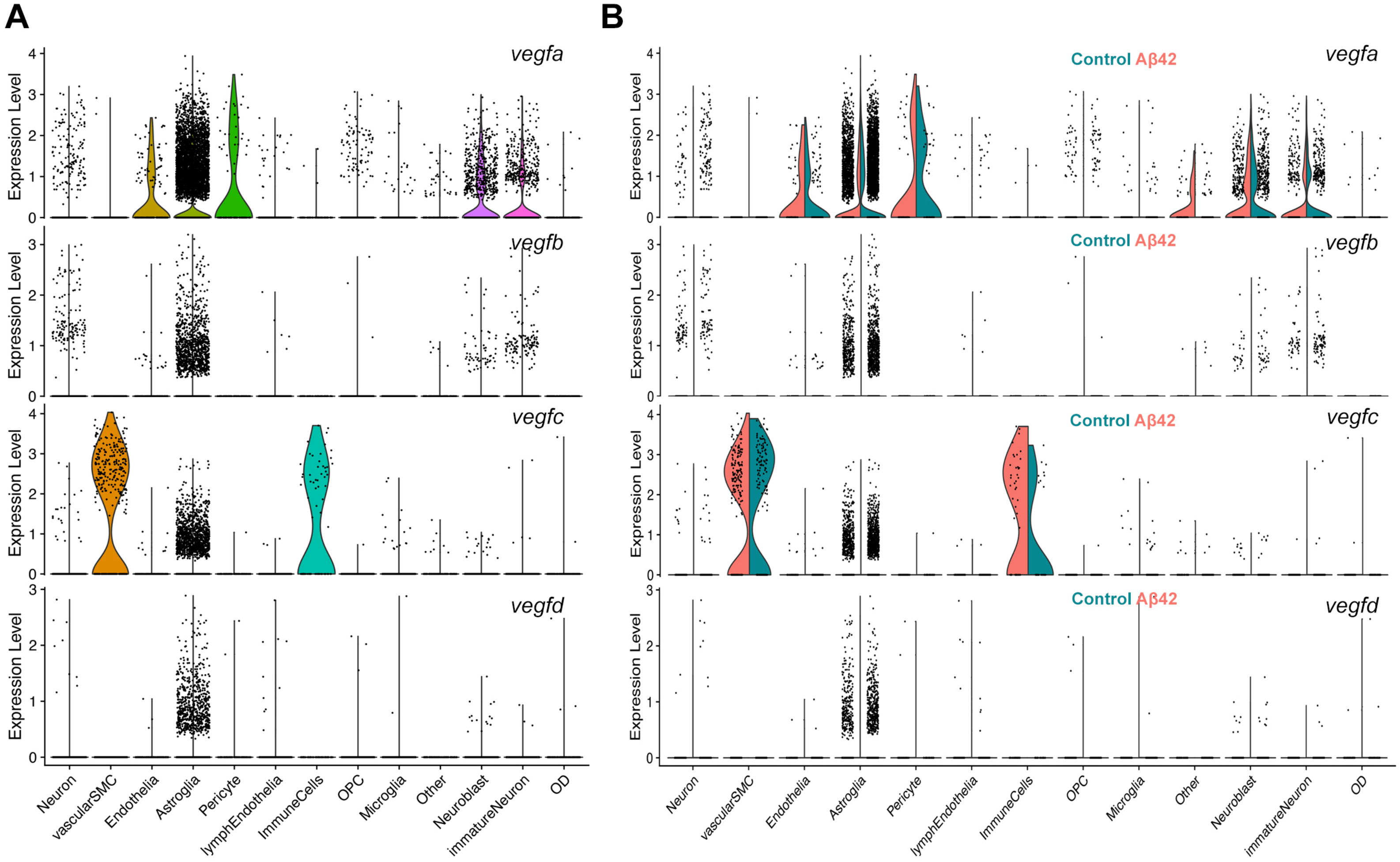

A
B
vegfa
vegfb
vegfc
vegfd
Control Aβ42
Expression Level
Neuron
vascularSMC
Endothelia
Astroglia
Pericyte
lymphEndothelia
ImmuneCells
OPC
Microglia
Other
Neuroblast
immatureNeuron
OD

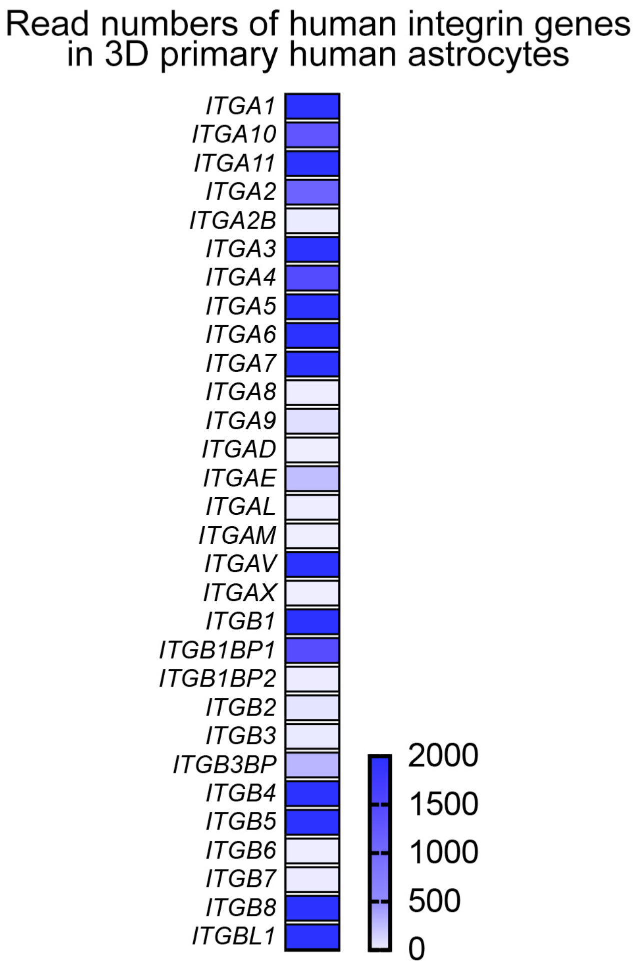
Read numbers of human integrin genes in 3D primary human astrocytes
ITGA1
ITGA10
ITGA11
ITGA2
ITGA2B
ITGA3
ITGA4
ITGA5
ITGA6
ITGA7
ITGA8
ITGA9
ITGAD
ITGAE
ITGAL
ITGAM
ITGAV
ITGAX
ITGB1
ITGB1BP1
ITGB1BP2
ITGB2
ITGB3
ITGB3BP
ITGB4
ITGB5
ITGB6
ITGB7
ITGB8
ITGBL1
2000
1500
1000
500
0

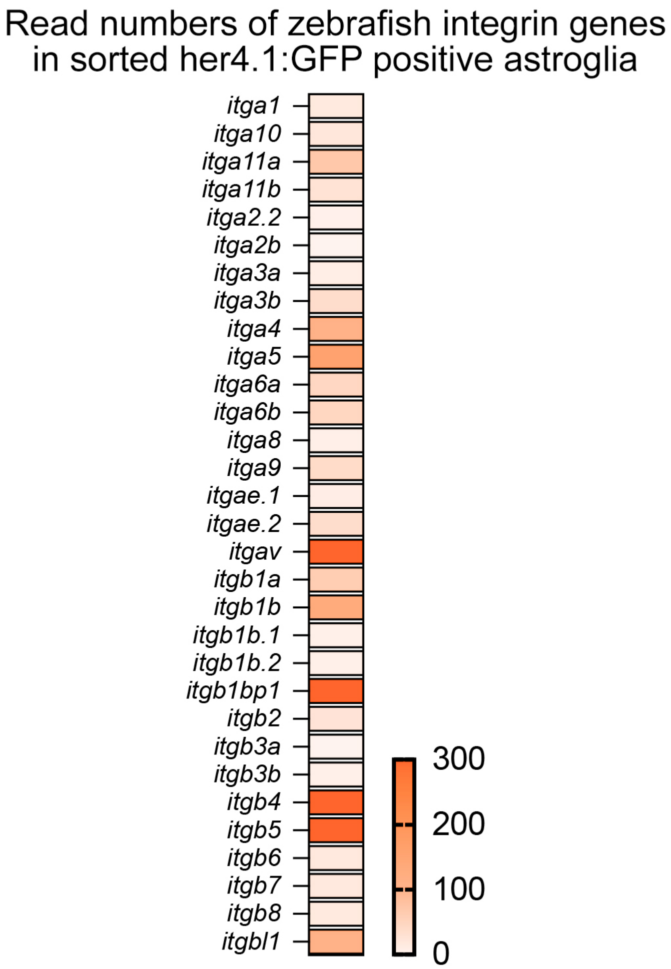
Read numbers of zebrafish integrin genes in sorted her4.1:GFP positive astroglia
itga1
itga10
itga11a
itga11b
itga2.2
itga2b
itga3a
itga3b
itga4
itga5
itga6a
itga6b
itga8
itga9
itgae.1
itgae.2
itgav
itgb1a
itgb1b
itgb1b.1
itgb1b.2
itgb1bp1
itgb2
itgb3a
itgb3b
itgb4
itgb5
itgb6
itgb7
itgb8
itgbl1
300
200
100
0

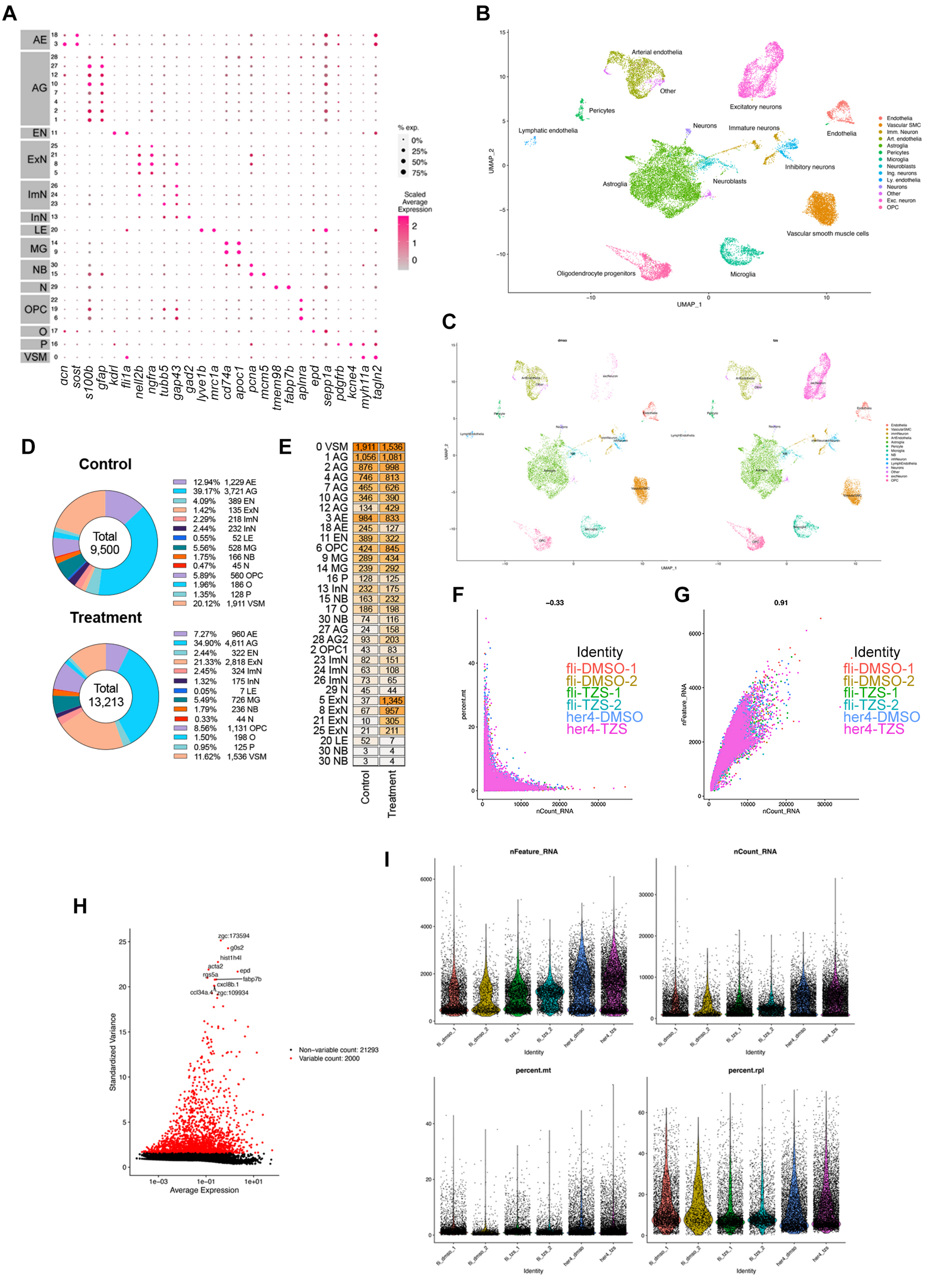
A
AE
AG
EN
ExN
ImN
InN
LE
MG
NB
N
OPC
O
P
VSM
acn
sost
s100b
gfap
kdrl
fli1a
nell2b
ngfra
tubb5
gap43
gad2
lyve1b
mrc1a
cd74a
apoc1
pcna
mcm5
tmem98
fabp7b
aplnra
epd
sepp1a
pdgfrb
kcne4
myh11a
tagln2
% exp.
0%
25%
50%
75%
Scaled Average Expression
B
Arterial endothelia
Other
Excitatory neurons
Pericytes
Lymphatic endothelia
Neurons
Immature neurons
Endothelia
Inhibitory neurons
Astroglia
Neuroblasts
Vascular smooth muscle cells
Oligodendrocyte progenitors
Microglia
UMAP_1
UMAP_2
Endothelia
Vascular SMC
Imm. Neuron
Art. endothelia
Astroglia
Pericytes
Microglia
Neuroblasts
Ing. neurons
Ly. endothelia
Neurons
Other
Exc. neuron
OPC
C
dmso
tzs
D
Control
Total 9,500
12.94% 1,229 AE
39.17% 3,721 AG
4.09% 389 EN
1.42% 135 ExN
2.29% 218 ImN
2.44% 232 InN
0.55% 52 LE
5.56% 528 MG
1.75% 166 NB
0.47% 45 N
5.89% 560 OPC
1.96% 186 O
1.35% 128 P
20.12% 1,911 VSM
Treatment
Total 13,213
7.27% 960 AE
34.90% 4,611 AG
2.44% 322 EN
21.33% 2,818 ExN
2.45% 324 ImN
1.32% 175 InN
0.05% 7 LE
5.49% 726 MG
1.79% 236 NB
0.33% 44 N
8.56% 1,131 OPC
1.50% 198 O
0.95% 125 P
11.62% 1,536 VSM
E
0 VSM 1,911 1,536
1 AG 1,056 1,081
2 AG 876 998
4 AG 746 813
7 AG 465 626
10 AG 346 390
12 AG 134 429
3 AE 984 833
18 AE 245 127
11 EN 389 322
6 OPC 424 845
9 MG 289 434
14 MG 239 292
16 P 128 125
13 InN 232 175
15 NB 163 232
17 O 186 198
30 NB 74 116
27 AG 24 158
28 AG2 93 203
2 OPC1 43 83
23 ImN 82 151
24 ImN 63 108
26 ImN 73 65
29 N 45 44
5 ExN 37 1,345
8 ExN 67 957
21 ExN 10 305
25 ExN 21 211
20 LE 52 7
30 NB 3 4
30 NB 3 4
Control
Treatment
F
−0.33
percent.mt
nCount_RNA
Identity
fli-DMSO-1
fli-DMSO-2
fli-TZS-1
fli-TZS-2
her4-DMSO
her4-TZS
G
0.91
nFeature_RNA
nCount_RNA
Identity
fli-DMSO-1
fli-DMSO-2
fli-TZS-1
fli-TZS-2
her4-DMSO
her4-TZS
H
zgc:173594
g0s2
hist1h4l
acta2
epd
rgs5a
fabp7b
cxcl8b.1
ccl34a.4
zgc:109934
Standardized Variance
Average Expression
Non-variable count: 21293
Variable count: 2000
I
nFeature_RNA
nCount_RNA
percent.mt
percent.rpl
fli_dmso_1
fli_dmso_2
fli_tzs_1
fli_tzs_2
her4_dmso
her4_tzs
Identity

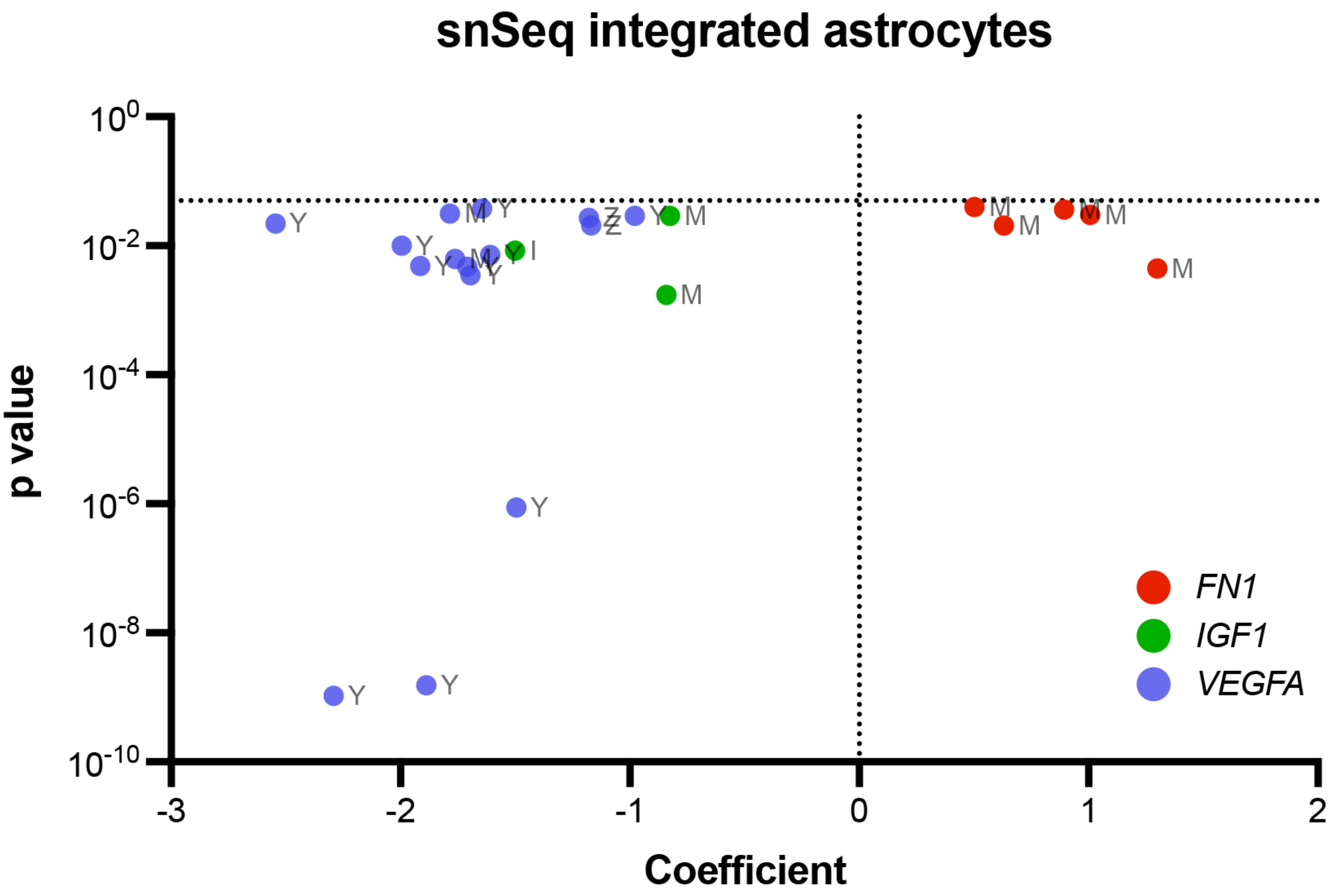
snSeq integrated astrocytes
p value
Coefficient
$10^0$
$10^{-2}$
$10^{-4}$
$10^{-6}$
$10^{-8}$
$10^{-10}$
-3
-2
-1
0
1
2
FN1
IGF1
VEGFA
Y
M
Z
I

### Electrophoresis File Run Summary

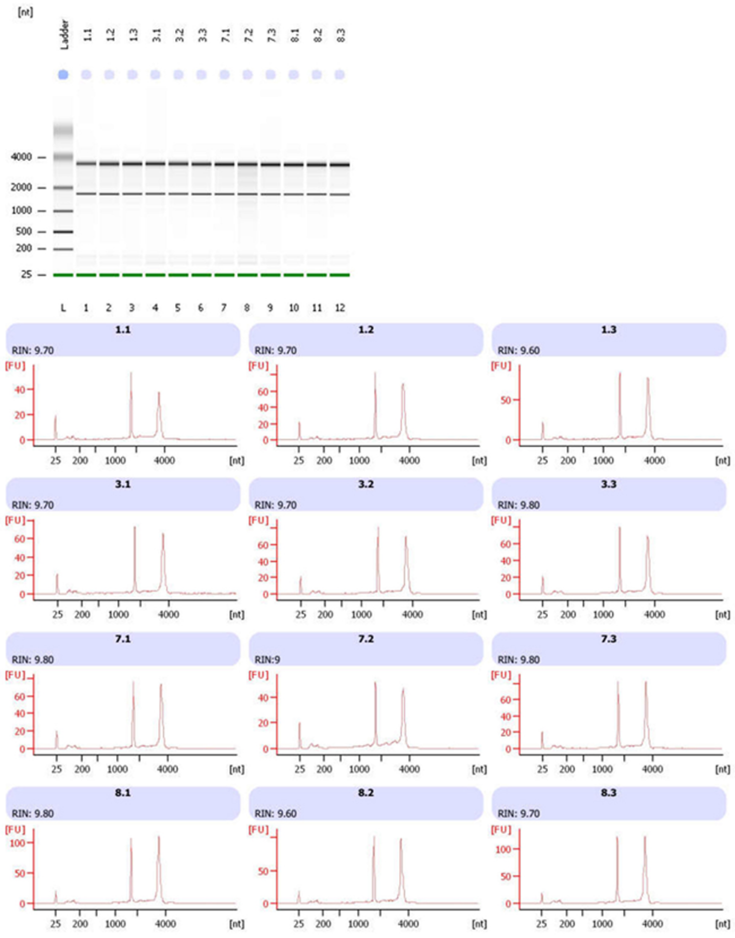


2100 Expert (B.02.08.SI648)

© Copyright 2003 - 2009 Agilent Technologies, Inc.

Printed: 2/23/2018 12:20:34 PM

### Electropherogram Summary

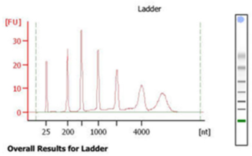


**Overall Results for Ladder**

| | |
|---|---|
| RNA Area: | 309.8 |
| RNA Concentration: | 150 ng/µl |
| Result Flagging Color: | |
| Result Flagging Label: | All Other Samples |

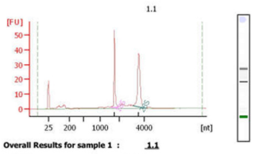


**Overall Results for sample 1 : 1.1**

| | |
|---|---|
| RNA Area: | 176.2 |
| RNA Concentration: | 85 ng/µl |
| rRNA Ratio [28s / 18s]: | 1.5 |
| RNA Integrity Number (RIN): | 9.7 (B.02.08) |
| Result Flagging Color: | |
| Result Flagging Label: | RIN: 9.70 |

**Fragment table for sample 1 : 1.1**

| Name | Start Size [nt] | End Size [nt] | Area | % of total Area |
|---|---|---|---|---|
| 18S | 1,534 | 1,930 | 43.6 | 24.8 |
| 28S | 3,106 | 4,074 | 63.5 | 36.1 |

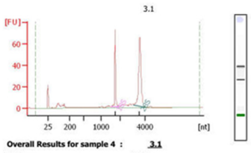


**Overall Results for sample 4 : 3.1**

| | |
|---|---|
| RNA Area: | 254.4 |
| RNA Concentration: | 123 ng/µl |
| rRNA Ratio [28s / 18s]: | 1.8 |
| RNA Integrity Number (RIN): | 9.7 (B.02.08) |
| Result Flagging Color: | |
| Result Flagging Label: | RIN: 9.70 |

**Fragment table for sample 4 : 3.1**

| Name | Start Size [nt] | End Size [nt] | Area | % of total Area |
|---|---|---|---|---|
| 18S | 1,524 | 1,955 | 58.5 | 23.0 |
| 28S | 3,065 | 4,066 | 104.2 | 41.0 |

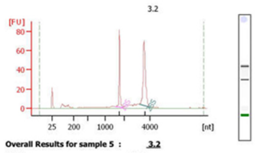


**Overall Results for sample 5 : 3.2**

| | |
|---|---|
| RNA Area: | 277.2 |
| RNA Concentration: | 134 ng/µl |
| rRNA Ratio [28s / 18s]: | 1.8 |
| RNA Integrity Number (RIN): | 9.7 (B.02.08) |
| Result Flagging Color: | |
| Result Flagging Label: | RIN: 9.70 |

**Fragment table for sample 5 : 3.2**

| Name | Start Size [nt] | End Size [nt] | Area | % of total Area |
|---|---|---|---|---|
| 18S | 1,508 | 1,947 | 63.8 | 23.0 |
| 28S | 3,130 | 4,062 | 112.5 | 40.6 |

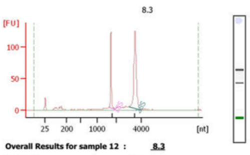


**Overall Results for sample 12 : 8.3**

| | |
|---|---|
| RNA Area: | 419.1 |
| RNA Concentration: | 203 ng/µl |
| rRNA Ratio [28s / 18s]: | 2.2 |
| RNA Integrity Number (RIN): | 9.7 (B.02.08) |
| Result Flagging Color: | |
| Result Flagging Label: | RIN: 9.70 |

**Fragment table for sample 12 : 8.3**

| Name | Start Size [nt] | End Size [nt] | Area | % of total Area |
|---|---|---|---|---|
| 18S | 1,644 | 1,897 | 91.1 | 21.7 |
| 28S | 3,007 | 3,996 | 196.9 | 47.0 |

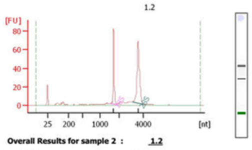


**Overall Results for sample 2 : 1.2**

| | |
|---|---|
| RNA Area: | 282.6 |
| RNA Concentration: | 137 ng/µl |
| rRNA Ratio [28s / 18s]: | 1.8 |
| RNA Integrity Number (RIN): | 9.7 (B.02.08) |
| Result Flagging Color: | |
| Result Flagging Label: | RIN: 9.70 |

**Fragment table for sample 2 : 1.2**

| Name | Start Size [nt] | End Size [nt] | Area | % of total Area |
|---|---|---|---|---|
| 18S | 1,514 | 1,942 | 66.9 | 23.7 |
| 28S | 3,073 | 4,006 | 120.6 | 42.7 |

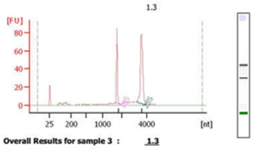


**Overall Results for sample 3 : 1.3**

| | |
|---|---|
| RNA Area: | 297.5 |
| RNA Concentration: | 144 ng/µl |
| rRNA Ratio [28s / 18s]: | 1.9 |
| RNA Integrity Number (RIN): | 9.6 (B.02.08) |
| Result Flagging Color: | |
| Result Flagging Label: | RIN: 9.60 |

**Fragment table for sample 3 : 1.3**

| Name | Start Size [nt] | End Size [nt] | Area | % of total Area |
|---|---|---|---|---|
| 18S | 1,676 | 1,936 | 63.4 | 21.3 |
| 28S | 3,148 | 4,006 | 122.9 | 41.3 |

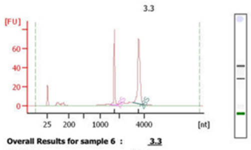


**Overall Results for sample 6 : 3.3**

| | |
|---|---|
| RNA Area: | 254.5 |
| RNA Concentration: | 123 ng/µl |
| rRNA Ratio [28s / 18s]: | 1.7 |
| RNA Integrity Number (RIN): | 9.8 (B.02.08) |
| Result Flagging Color: | |
| Result Flagging Label: | RIN: 9.80 |

**Fragment table for sample 6 : 3.3**

| Name | Start Size [nt] | End Size [nt] | Area | % of total Area |
|---|---|---|---|---|
| 18S | 1,521 | 1,936 | 63.1 | 24.8 |
| 28S | 3,068 | 3,952 | 110.4 | 43.4 |

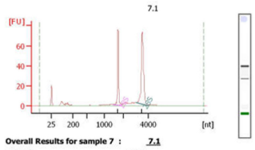


**Overall Results for sample 7 : 7.1**

| | |
|---|---|
| RNA Area: | 261.5 |
| RNA Concentration: | 127 ng/µl |
| rRNA Ratio [28s / 18s]: | 1.9 |
| RNA Integrity Number (RIN): | 9.8 (B.02.08) |
| Result Flagging Color: | |
| Result Flagging Label: | RIN: 9.80 |

**Fragment table for sample 7 : 7.1**

| Name | Start Size [nt] | End Size [nt] | Area | % of total Area |
|---|---|---|---|---|
| 18S | 1,493 | 1,935 | 60.2 | 23.0 |
| 28S | 3,029 | 3,996 | 114.7 | 43.9 |

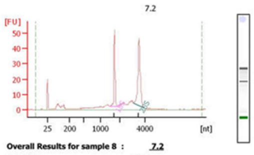


**Overall Results for sample 8 : 7.2**

| | |
|---|---|
| RNA Area: | 242.8 |
| RNA Concentration: | 118 ng/µl |
| rRNA Ratio [28s / 18s]: | 1.7 |
| RNA Integrity Number (RIN): | 9 (B.02.08) |
| Result Flagging Color: | |
| Result Flagging Label: | RIN:9 |

**Fragment table for sample 8 : 7.2**

| Name | Start Size [nt] | End Size [nt] | Area | % of total Area |
|---|---|---|---|---|
| 18S | 1,481 | 1,908 | 42.9 | 17.7 |
| 28S | 3,085 | 3,922 | 73.6 | 30.3 |

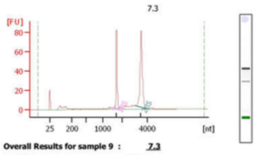


**Overall Results for sample 9 : 7.3**

| | |
|---|---|
| RNA Area: | 292.1 |
| RNA Concentration: | 141 ng/µl |
| rRNA Ratio [28s / 18s]: | 1.9 |
| RNA Integrity Number (RIN): | 9.8 (B.02.08) |
| Result Flagging Color: | |
| Result Flagging Label: | RIN: 9.80 |

**Fragment table for sample 9 : 7.3**

| Name | Start Size [nt] | End Size [nt] | Area | % of total Area |
|---|---|---|---|---|
| 18S | 1,489 | 1,925 | 65.5 | 22.4 |
| 28S | 3,018 | 3,989 | 124.8 | 42.7 |

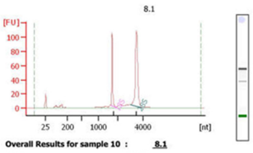


**Overall Results for sample 10 : 8.1**

| | |
|---|---|
| RNA Area: | 371.5 |
| RNA Concentration: | 180 ng/µl |
| rRNA Ratio [28s / 18s]: | 2.1 |
| RNA Integrity Number (RIN): | 9.8 (B.02.08) |
| Result Flagging Color: | |
| Result Flagging Label: | RIN: 9.80 |

**Fragment table for sample 10 : 8.1**

| Name | Start Size [nt] | End Size [nt] | Area | % of total Area |
|---|---|---|---|---|
| 18S | 1,651 | 1,897 | 78.9 | 21.2 |
| 28S | 3,035 | 3,972 | 168.8 | 45.4 |

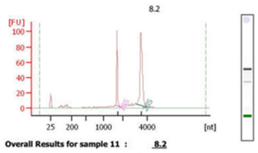


**Overall Results for sample 11 : 8.2**

| | |
|---|---|
| RNA Area: | 349.6 |
| RNA Concentration: | 169 ng/µl |
| rRNA Ratio [28s / 18s]: | 2.1 |
| RNA Integrity Number (RIN): | 9.6 (B.02.08) |
| Result Flagging Color: | |
| Result Flagging Label: | RIN: 9.60 |

**Fragment table for sample 11 : 8.2**

| Name | Start Size [nt] | End Size [nt] | Area | % of total Area |
|---|---|---|---|---|
| 18S | 1,650 | 1,905 | 73.4 | 21.0 |
| 28S | 3,077 | 3,962 | 152.5 | 43.6 |
